# A morphometric double dissociation: cortical thickness is more related to aging; surface area is more related to cognition

**DOI:** 10.1101/2021.09.30.462545

**Authors:** G. Sophia Borgeest, Richard N. Henson, Tim C. Kietzmann, Christopher R. Madan, Theresa Fox, Maura Malpetti, Delia Fuhrmann, Ethan Knights, Johan D. Carlin, Cam-CAN, Rogier A. Kievit

## Abstract

The *thickness* and *surface area* of cortex are genetically distinct aspects of brain structure, and may be affected differently by age. However, their potential to differentially predict age and cognitive abilities has been largely overlooked, likely because they are typically aggregated into the commonly used measure of *volume*. In a large sample of healthy adults (N=647, aged 18-88), we investigated the brain-age and brain-cognition relationships of thickness, surface area, and volume, plus five additional morphological shape metrics. Cortical thickness was the metric most strongly associated with age cross-sectionally, as well as exhibiting the steepest longitudinal change over time (subsample N=261, aged 25-84). In contrast, surface area was the best single predictor of age-residualized cognitive abilities (fluid intelligence), and changes in surface area were most strongly associated with cognitive change over time. These findings were replicated in an independent dataset (N=1345, aged 18-93). Our results suggest that cortical thickness and surface area make complementary contributions the age-brain-cognition triangle, and highlight the importance of considering these volumetric components separately.

## Introduction

As the human brain ages, it undergoes a pronounced structural transformation. Even in the absence of neuropathology, overall brain volume shrinks – from age six onwards into old age (Bethlehem et al., 2021). This volume decline is associated with various physiological changes, including grey-matter reductions caused largely by the regression of dendrites (see Dickstein et al., 2007 for a review), and white-matter reductions stemming from axon demyelination (Fotenos et al., 2005; Gunning-Dixon et al., 2009; Raz, 2005; Scheltens et al., 1995). There are also morphological changes, with sulci for example becoming shallower (Burgmans et al., 2011; Jin et al., 2018; Madan, 2021; Peters, 2007) and cortex becoming more curved (Deppe et al., 2014).

Traditionally, studies investigating human brain structure with Magnetic Resonance Imaging (MRI) have relied largely on volumetric or thickness measures (see Oschwald et al., 2020 for a review), which only capture a small proportion of the richness of age-related morphometric changes (Ecker et al., 2010; Im et al., 2008). Indeed, the number of papers that include both the term “aging” and “brain volume” (N=2715 in a PubMed search as of 01/06/2021) or “cortical thickness” (N=597) far exceeds those investigating other aspects of morphology, such as “aging” combined with “surface area” (N=125) or “curvature” (N=23). Even though several authors have pointed out that volume is a product of cortical thickness and surface area (Norbom et al., 2021; Storsve et al., 2014; Walhovd et al., 2016; Winkler et al., 2018), which in turn are two genetically independent aspects of brain structure (Hofer et al., 2020; McKay et al., 2014; Panizzon et al., 2009; van der Meer et al., 2020), the implication that thickness and area may have dissociable causes (e.g., in ageing) and consequences (e.g., for cognition) have rarely been discussed, especially in adult samples. Moreover, additional detailed morphometric shape measures (such as curvature or sulcal depth) may provide further insight into brain development across the adult lifespan and its relationship with cognitive performance.

In this paper, we explore multiple morphometric measures in two large adult-lifespan cohorts. We show, firstly, that the most pronounced structural changes in the aging brain are the decrease in apparent cortical thickness (see Walhovd et al., 2017 for the interpreation of MR-derived cortical thickness) and increase in cortical curvature, in line with other studies (Deppe et al., 2014; Hogstrom et al., 2013; Lemaitre et al., 2012). Secondly, we find that incorporating multiple shape measures into a single model outperforms any individual metrics’ ability to capture age-related and fluid cognitive differences. This paper’s main contribution, however, lies in providing cross-sectional and longitudinal evidence of a double dissociation in two independent, large-sample cohorts. Specifically, cortical thickness was more strongly associated with age than cortical surface area, while surface area was more strongly associated with cognition (as indexed by fluid intelligence). This pattern was most apparent longitudinally, but we also observed it cross-sectionally after adjusting for age. This double dissociation points to possibly distinct underlying biological processes (discussed below), and supports recent calls to investigate thickness and surface area separately (Winkler et al., 2018) as brain volume (a product of cortical thickness and surface area) likely conflates and therefore masks these differentiable effects.

## Results

### Cross-sectional results

We first calculated whole brain as well as regional correlations between each metric and age, cognitive abilities (as indexed by fluid intelligence) and age-residualized cognitive abilities Residualized cognitive scores allow one to separate concurrent age-related decline in cognitive ability, thus providing an age-independent measure of cognition. Thickinthehead, which is a measure of cortical thickness from the Mindboggle software, showed the strongest whole-brain-age correlations (r = −.83). This was followed by curvature (r = +.77), fractal dimensionality (a measure of cortical complexity; = −.65) and FreeSurfer’s standard cortical thickness (r = −.60), as shown in Table 1 and plotted in Figure 1. Compared to the other metrics, surface area exhibited the weakest age relationship (r = −.36). This order was reversed for age-residualized cognition. Here, surface area was the strongest predictor (r = +0.21), while the two thickness metrics and curvature did not show significant brain-cognition correlations after adjusting for age. The two volume measures (FreeSurfer’s cortical volume, plus SPM’s cortical + subcortical volume) predicted both age and age-residualized fluid-intelligence reasonably well (r ~ −.55 and 0.20, respectively), as would be expected since they are proportional to the product of cortical thickness and surface area. Fractal dimensionality was also a good predictor of both age and age-residualized cognition (r_age_= −0.65, r_cog_ = 0.19).

**Table 1:** whole brain correlations. GM = grey-matter. FS = FreeSurfer. SPM = Statistical Parametric Mapping. MB = Mindboggle.

| Metric | Age |  | Fluid Intelligence |  | Age-residualized Fluid Intelligence |  |
| --- | --- | --- | --- | --- | --- | --- |
|  | Pearson's | *P* | Pearson's | *p* | Pearson's | *P* |
|  | *r* |  | *r* |  | *r* |  |
| Cortical Volume (FS) | -.62 | <.001 | +.56 | <.001 | +.20 | <.001 |
| Cortical Thickness (FS) | -.60 | <.001 | +.42 | <.001 | +.04 | .33 |
| Surface Area (FS) | -.36 | <.001 | +.39 | <.001 | +.21 | <.001 |
| Thickinthehead (MB) | -.83 | <.001 | +.59 | <.001 | +.04 | .34 |
| Curvature (MB) | +.77 | <.001 | -.56 | <.001 | -.034 | .39 |
| Sulcal Depth (MB) | -.38 | <.001 | +.51 | <.001 | +.07 | .06 |
| GM Volume (SPM) | -.54 | <.001 | +.51 | <.001 | +.20 | <.001 |
| Fractal Dimensionality | -.65 | <.001 | +.56 | <.001 | +.19 | <.001 |

**Figure 1:**
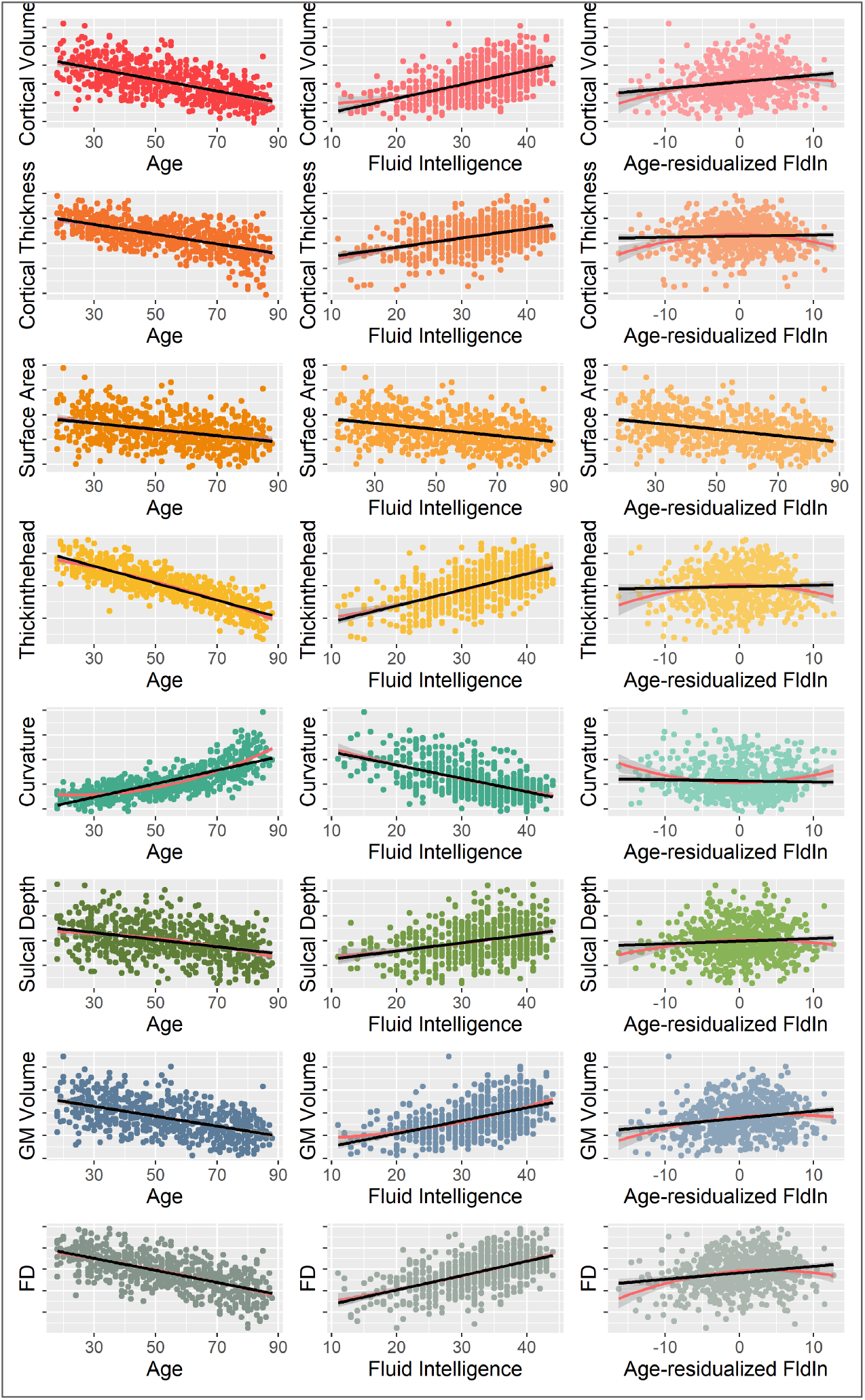
whole brain -age, -fluid intelligence and -age-residualized fluid intelligence scatterplots of all eight metrics. Black lines show linear fit, red lines show quadratic fit. The metric exhibiting the strongest age relationship is Thickinthehead (a measure of cortical thickness), while surface area is most strongly related to age-residualized cognitive abilities. GM = Grey Matter, FD = Fractal Dimensionality.

**Figure 2:**
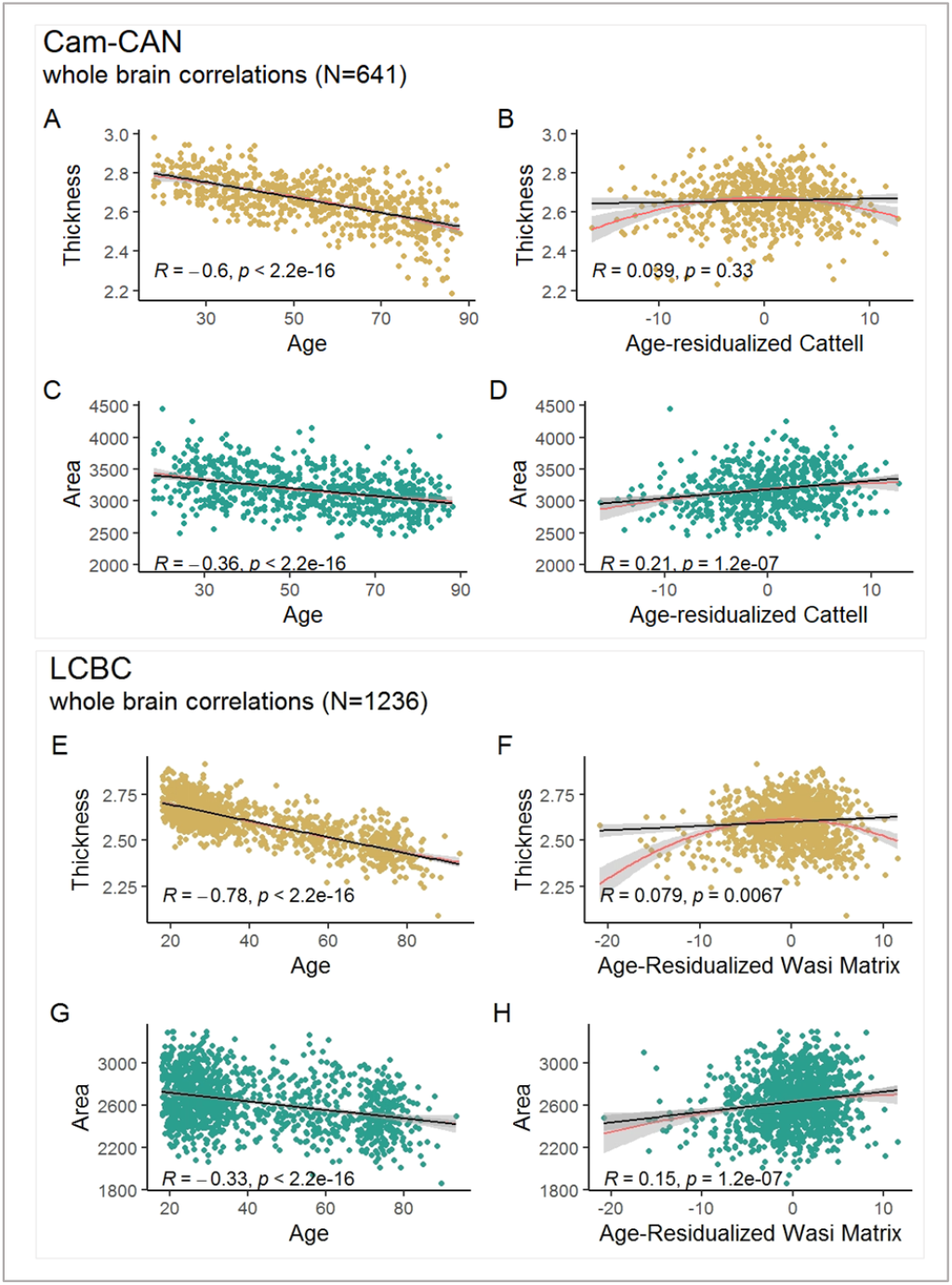
Cross-sectional whole brain correlations in Cam-CAN (A-D) and LCBC (E-H). While thickness is associated with age (not age-residualized cognition), surface area captures age-residualized cognition well (and age comparatively poorly).

Next, we estimated a series of path models to assess the relationship between brain structure and age, fluid intelligence and age-residualized fluid intelligence when both surface area and cortical thickness are included in the same model. Path analysis is an extension of multiple linear regressions, allowing researchers to assess the relationships between the predictor variables rather than having several independent variables predict one dependent variable (Streiner, 2005). Age and fluid intelligence were best captured by surface area and cortical thickness, while age-residualized fluid intelligence was associated only with surface area (see Figure 3). We validated this frequentist modelling approach with Bayesian model selection (supplementary Figures 4–5). Overall, the whole-brain, cross-sectional analyses suggest that cortical thickness and surface area differentially associated with age and age-residualized cognitive abilities, respectively.

**Figure 3:**
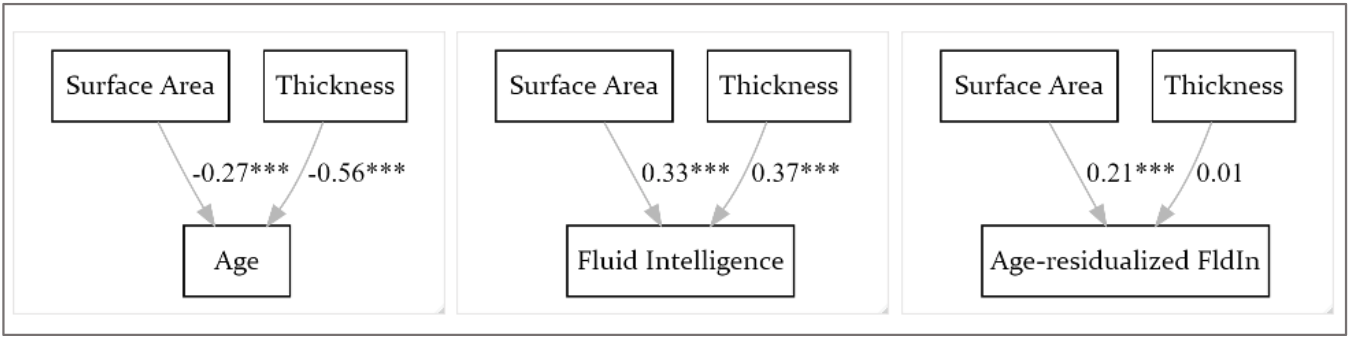
Cam-CAN path model results. Both surface area and thickness are significantly associated with age and fluid intelligence, while age-residualized fluid intelligence is captured by surface area only.

Our regional investigations further support the morphological dichotomy found in the whole brain analyses. As shown in Figure 4, for cortical thickness, all 32 brain regions (the 64 DKT regions averaged across the hemispheres) were significantly correlated with age (all correlations were FDR corrected at alpha = 0.05), while no region predicted age-residualized fluid intelligence (r < 0.07, P_FDR_ > 0.05; see supplementary tables 5–7). In contrast, for surface area, *all* regions were significantly associated with age-residualized cognitive abilities (r > 0.11, P_FDR_ < 0.05). While regional surface area also correlated with age, the correlations were substantially weaker than the brain-age correlations for cortical thickness.

**Figure 4:**
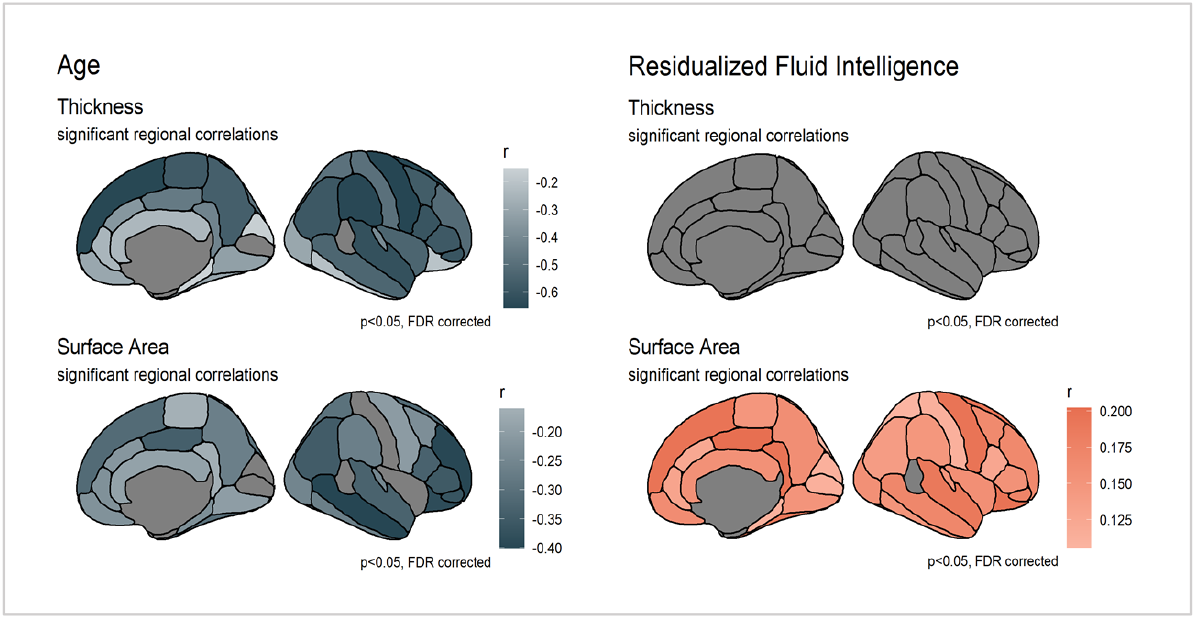
Significant regional age- and age-residualized fluid intelligence correlations. Correlations are FDR corrected at alpha = 0.05. For cortical thickness, all 32 brain regions are significantly associated with age, while none are associated with age-residualized cognitive abilities. For surface area, all regions are correlated with age-residualized cognition. While regional surface area also correlated with age, the correlations were substantially weaker than the brain-age correlations for cortical thickness.

**Figure 5:**
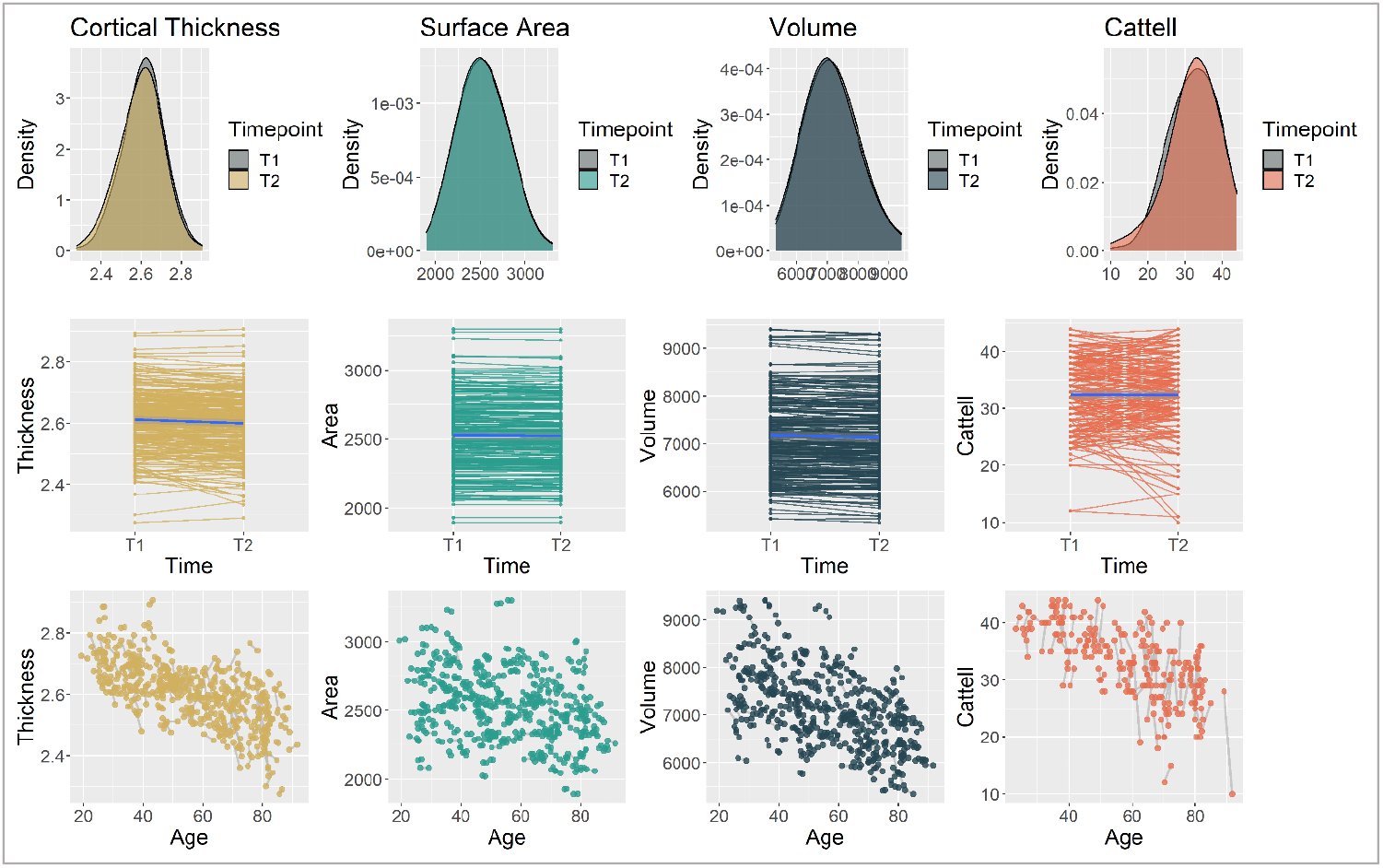
In Cam-CAN, cortical thickness, surface area and fluid intelligence declined significantly between time point 1 and time point 2 (average interval between the two time points = 1.33 years).

**Figure 6:**
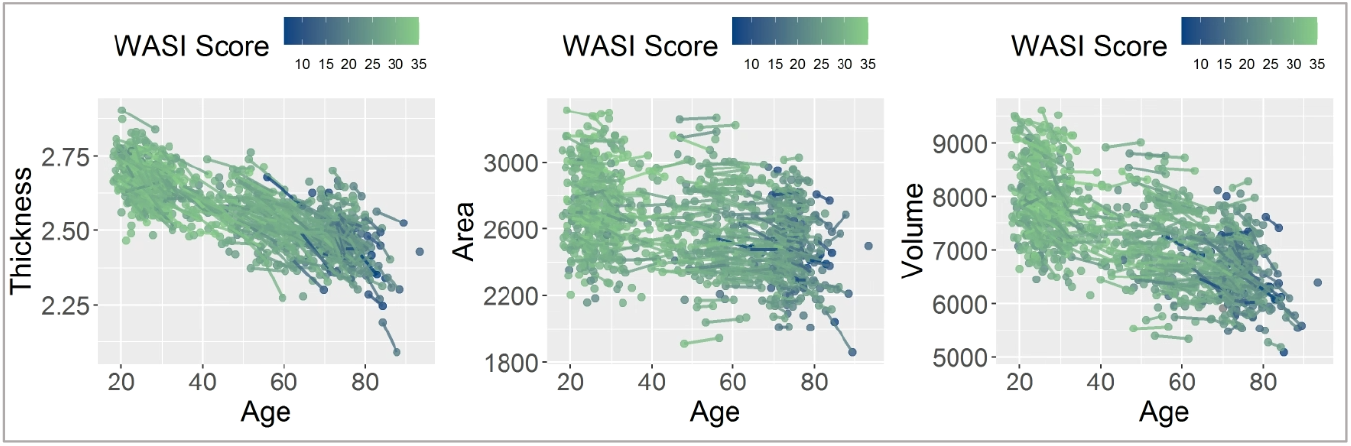
The relationship between age, brain structure and cognition in LCBC.

Finally, in addition to the “area and thickness only” path models, we ran three “full models” which each included all eight brain structure metrics to assess the metrics’ combined associations with age and cognition. The total variance explained by these models was 76, 46 and 7 percent for age, fluid intelligence and age-residualized fluid intelligence, respectively – almost double the variance explained by thickness and area alone (see supplementary Figure 3). Moreover, the fact that multiple morphometric measures provided partially complementary information about the outcome highlights the potential usefulness in assessing various morphological shape measures when investigating the ageing brain and cognitive abilities. This was further supported by regional brain-age and brain-cognition correlations (supplementary Figure 8): for instance, while volume-age effects were most pronounced in the frontal regions, depth-age effects were strongest in the temporal lobes. It is plausible that the focus on frontal brain regions in the brain and cognitive aging literature (Greenwood, 2000; Jung & Haier, 2007) is informed in part by the field’s traditional focus on brain volume, and that other aspects of brain structure could point to more underappreciated regional effects.

### Longitudinal results

Although cross-sectional analyses offer an interesting insight into age-related cognitive and morphometric *differences*, longitudinal data are needed to truly assess how brain and cognitive *change* (Oschwald et al., 2020). Doing so, we found that the change-change relationship between surface area and cognition was significantly stronger than the change-change relationship between volume and cognition as well as that between thickness and cognition.

After establishing metric and scalar invariance (described in supplementary section 7), we used Latent Change Score Models (LCSM) to examine morphometric and cognitive change over time. The cognitive LCSM revealed significant change in cognition over time, as well as significant variability in the rate of change (Table 2, variances). The effect size of change of fluid intelligence was −0.04 (Cohen’s D, computed by dividing the mean change by the SD at time 1). The three brain-structure LCSMs also showed evidence of change over time (Table 2, intercepts) and of significant variability in the rate of change (Table 2, variances). Surface area, volume and thickness all decreased between the first and the second scan. Surface area had the smallest effect size (Cohen’s D = −0.02), with cortical thickness and volume exhibiting larger effects (Cohen’s D of −0.12 and −0.11, respectively).

**Table 2:** latent change score model results for change in Cattell, surface area, thickness and volume over time. Effect size is calculated by dividing the mean change by the square root of the variance.

|  |  | Estimate | SE | z-value | p | Std.all | Effect size |
| --- | --- | --- | --- | --- | --- | --- | --- |
| **Cattell** | Intercepts | -0.633 | 0.289 | -2.192 | 0.028 | -0.145 | -0.09 |
|  | Variances | 19.059 | 2.808 | 6.787 | <.0001 | 1.000 |  |
| **Thickness** | Intercepts | -0.012 | 0.002 | -6.234 | <.0001 | -0.386 | -0.12 |
|  | Variances | 0.001 | 0.000 | 7.229 | <.0001 | 1.000 |  |
| **Surface Area** | Intercepts | -5.680 | 1.632 | -3.481 | <.0001 | -0.215 | -0.02 |
|  | Variances | 695.026 | 197.495 | 3.519 | <.0001 | 1.000 |  |
| **Volume** | Intercepts | -50.550 | 5.887 | -8.587 | <.0001 | -0.530 | -0.11 |
|  | Variances | 9080.25 | 1057.968 | 8.587 | <.0001 | 1.000 |  |

Next, to investigate the relationship between cognitive change and morphometric change, we fit three second order latent change score models (2LCSM), one for each brain structure metric. We used full information maximum likelihood (FIML, Enders & Mansolf, 2018) with robust standard errors to account for missing data. Results are shown in Table 3.

**Table 3:**
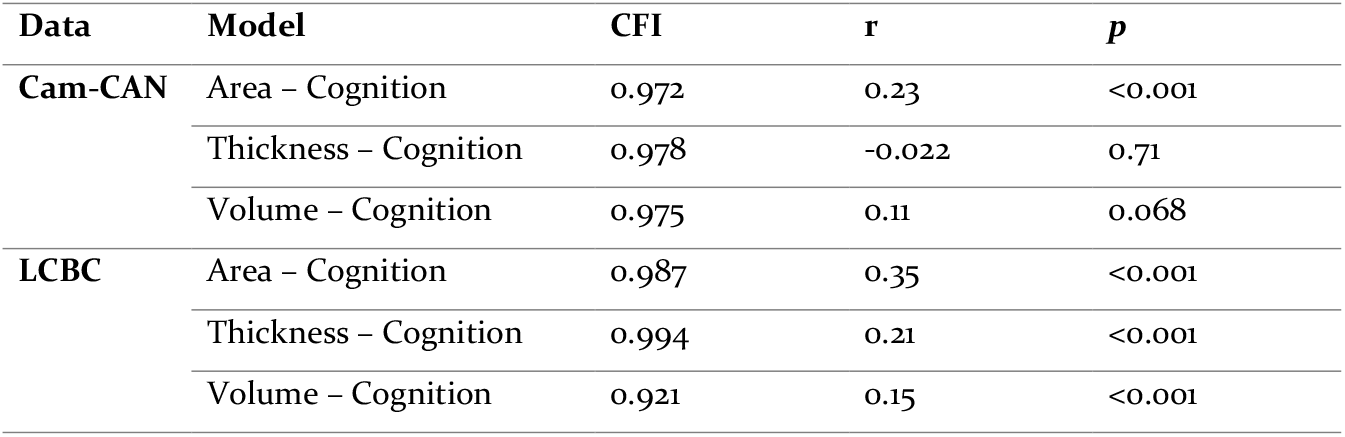
Second order latent change score model results using FIML for missing data. Shows the relationship between change in brain structure (volume, thickness, area) and change in cognition in Cam-CAN and LCBC. In both datasets, change in surface area was most strongly associated with cognitive change.

| Data | Model | CFI | r | p |
| --- | --- | --- | --- | --- |
| Cam-CAN | Area – Cognition | 0.972 | 0.23 | <0.001 |
|  | Thickness – Cognition | 0.978 | -0.022 | 0.71 |
|  | Volume – Cognition | 0.975 | 0.11 | 0.068 |
| LCBC | Area – Cognition | 0.987 | 0.35 | <0.001 |
|  | Thickness – Cognition | 0.994 | 0.21 | <0.001 |
|  | Volume – Cognition | 0.921 | 0.15 | <0.001 |

All three models fit the data well: CFI _area_ = 0.972; CFI _volume_ = 0.975; CFI _thickness_ = 0.978; (further model fit indices can be found in section 7 of the supplementary materials). After fitting the models, we extracted and correlated the cognitive rates of change with the brain structural rates of change. Change in surface area showed the largest effect (r = 0.23, p <.001), followed by (non-significantly) volume (r=-0.11, p = 0.068) and cortical thickness (r=-0.022, p = 0.71). The Steiger’s-Z tests (Steiger, 1980) in the R package psych can directly compare differences in correlation strengths, accounting for the full correlation pattern among variables. Doing so revealed that change in area was significantly more strongly associated with change in cognition than was thickness or volume change (see Table 4).

**Table 4:** Steiger’s Z Test results. *P*-value (two-tailed) of <0.05 suggests correlation coefficients are significantly different from each other.

| Data | Comparison | r values | N | Z | p |
| --- | --- | --- | --- | --- | --- |
| Cam-CAN | Thickness / Area | -0.022/0.23 | 362 | 3.34 | 0.001 |
|  | Thickness / Volume | -0.022/0.11 | 362 | 1.66 | 0.1 |
|  | Volume / Area | 0.11/0.23 | 362 | 1.77 | 0.04 |
| LCBC | Thickness / Area | 0.21/0.35 | 722 | 2.89 | 0.001 |
|  | Thickness / Volume | 0.21/0.15 | 722 | 1.18 | 0.24 |
|  | Volume / Area | 0.15/0.35 | 722 | 4.06 | 0.001 |

These results suggest that people whose surface area decreased more quickly also showed steeper rates of cognitive decline; an effect not found for thickness or volume.

Note that the models shown above include observed (not latent) variables to ensure maximum comparability between the LCBC and Cam-CAN models (in LCBC, it was not possible to derive latent cognitive scores because only WASI sum scores were available). However, latent variable Cam-CAN models (which we had run initially, before the replication study) show the same pattern, with changes in surface area most strongly associated with changes in cognition (r=0.44, p <0.001). For these models, changes in volume were significantly associated with changes in fluid intelligence (r=0.26, p = <0.001), while this relationship remained insignificant for cortical thickness (r = 0.0047, *p* = 0.94). All longitudinal change score model results are plotted in supplementary Figure 13.

### Replication results

To examine whether our cross-sectional and longitudinal findings generalize to other cohorts, we next (after finalizing the analyses in Cam-CAN) examined the same associations in an independent sample, the LCBC data. Because of their widespread use and accessibility, we included the three FreeSurfer-derived metrics (thickness, area, volume) in our replication analyses.

Cross-sectionally, as shown in Figure 2 (E-H), thickness showed the strongest whole brain-age correlation (R = −.78, *p* < 0.001), followed by volume (R = −0.64, *p* < 0.001) then surface area (R = −0.34, *p* < 0.001). For age-residualized fluid intelligence, thickness had the weakest correlation (R = 0.077, *p* = 0.009), followed by surface area (R = 0.13, *p* = 0.001) and volume (0.15, *p* < 0.001; and supplemental Table 3). As was the case in Cam-CAN, the frequentist path models and Bayesian model selection revealed that the best models to predict age and fluid intelligence were comprised of both surface area and thickness, while age-residualized fluid intelligence was best captured by surface area alone (Figure *7*).

**Figure 7:**
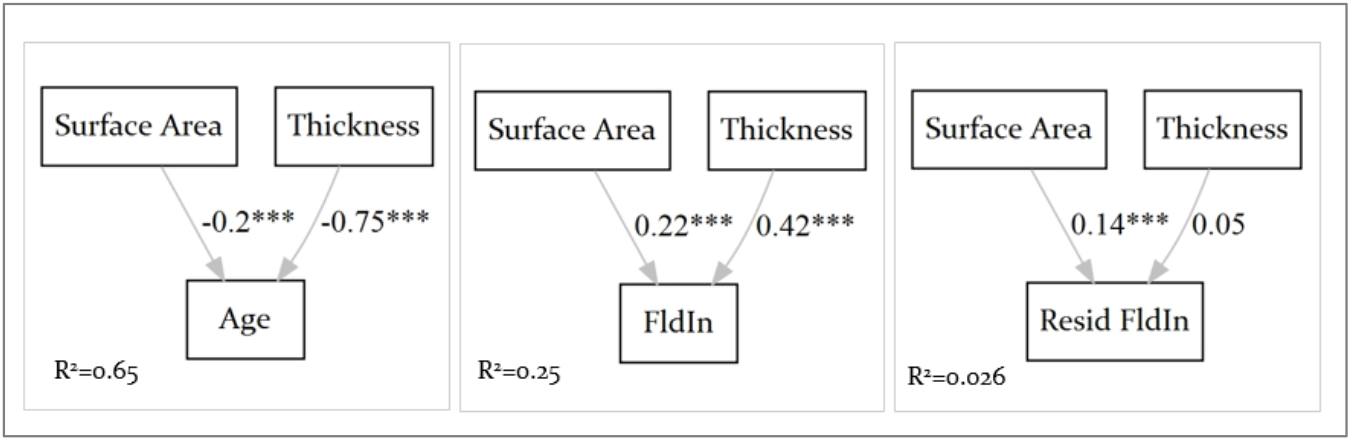
LCBC path model results. Both surface area and thickness are significantly associated with age and fluid intelligence, while age-residualized fluid intelligence is captured by surface area only.

Longitudinally, we found evidence of significant change over time for the three brain metrics (Table 5, intercepts), and significant variability over time for the brain metrics and cognition (Table 5, variances). A lack of mean cognitive decline can most likely be attributed to test-retest effects, but still allows for investigation of individual differences in change.

**Table 5:**
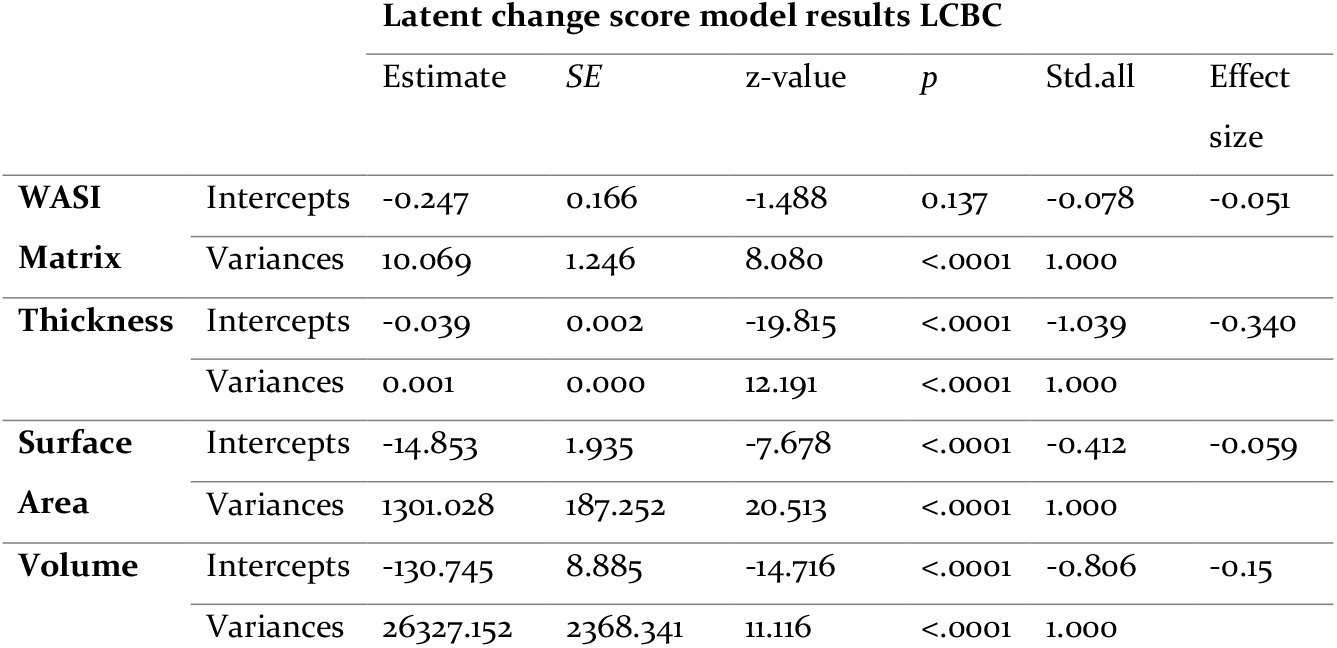
LCBC data latent change score model results for change in WASI Matrix, surface area, thickness and volume over time. Effect size is calculated by dividing the mean change by the square root of the variance.

As shown in Table 3, the three 2LCMs fit the data well: CFI area = 0. 0.987; CFI volume = 0.921; CFI thickness = 0.994 (further model fit indices can be found in the supplementary materials). Change in all structural brain metrics was significantly associated with change in cognition with surface area showing the largest effect (r = 0.35, *p* <.001), followed by thickness (r=0.22, *p* <.001) then volume (r=0.15, *p* =0.001). The Steiger’s Z-Test revealed that the change-change relationship between area and cognition was significantly stronger than that between volume and cognition and thickness and cognition (see Table 4).

The LCBC longitudinal results replicated those found in Cam-CAN, further supporting the finding that changes in surface area predict changes in cognition and that this relationship is stronger than that between change in thickness and change in cognition. We therefore successfully replicated Cam-CAN’s cross-sectional and longitudinal findings.

## Discussion

### A morphometric double dissociation

Across two independent cohorts, we found evidence of a morphometric double dissociation: cortical thickness was more strongly associated with age than cortical surface area, both cross-sectionally and longitudinally, whereas surface area was more strongly associated with cognition (fluid intelligence); certainly longitudinally, and also cross-sectionally, after removing age-related variance. Note that we are not claiming that cortical thickness plays *no* role in cognition – it shows a longitudinal association with cognitive change in one of the two datasets (albeit significantly smaller than that of surface area), and its cross-sectional association with fluid intelligence was significant. The lack of cross-sectional association with age-residualized fluid intelligence could be due to collider bias whereby cortical thickness is causally related to both age and cognition and that any thickness-cognition effect disappears when removing age. Our results do suggest, however, that surface area and thickness, which tend to be investigated together through the aggregate measure of volume, may have dissociable causes (e.g., in ageing) and consequences (e.g., for cognition).

Our findings align with previous studies that have pointed to a relationship between surface area and cognition (Cox et al., 2018; Fjell et al., 2015; Gerrits et al., 2016) and support recent calls to focus on the distinctness of cortical thickness and surface area, rather than assessing them jointly through cortical volume (Winkler et al., 2018). Such a shift is not just of theoretical or methodological importance: because surface area and cortical thickness are known to be genetically distinct (Panizzon et al., 2009; Winkler et al., 2010) and to follow different trajectories over the lifespan (Fjell et al., 2015; Hogstrom et al., 2013), combining them into volume is likely to obscure important biological differences and mechanisms.

While we can, in the present study, only speculate on the biological basis of different morphological metrics (and therefore their age/cognition dichotomy), evidence from animal and histological studies point to a possibly relevant set of mechanisms. With age, the long dendrites of pyramidal neurons have been shown to decrease rapidly across all layers of the cortex (Jacobs et al., 2001; Nakamura et al., 1985; Panizzon et al., 2009) and especially in layer V – the internal pyramidal layer – which contains the majority of large pyramidal neurons and is therefore the thickest of the six cortical layers – at least after the age of 50 (de Brabander et al., 1998). Thus, the steep declines in cortical thickness observed in the present study (and elsewhere, e.g. Lemaitre et al., 2012; Chen et al., 2011) are likely in part due to dendritic shrinkage. Furthermore, our finding that cortical thickness is less strongly associated with cognitive abilities than other measures of brain structure is also supported by animal research, showing that rates of dendritic atrophy in rats did not differ between aged cognitive unpaired and aged cognitive impaired animals (Allard et al., 2012)

What, if not dendritic atrophy, is driving cognitive differences and cognitive change, and why might cognition be related to surface area? According to the radial unit hypothesis (Rakic, 2000) while the development of cortical thickness is driven by the layers in the cortical columns (as described above), the development of surface area is a product of the number of radial columns perpendicular to the pial surface. This theory has been updated via the Supragranular Cortex Expansion Hypothesis (Nowakowski et al., 2016), which postulates that specific cellular mechanisms allow certain types of glial cells to migrate towards the pial surface during development, thereby expanding the cortex, and that this process is, in turn, responsible for many of the cognitive features unique to primates. This is further supported by analyses suggesting that glial cells – and specifically glial-neural signalling – affect cognition (Chung et al., 2015). A plausible hypothesis therefore is that MR-derived surface area (at least partially) picks up on these glial-dependent neural mechanisms – which likely originate in early development – and thereby on cognitive difference and changes.

### The shape of the ageing brain

A second contribution this paper makes is to characterize structural age-related differences and changes across multiple morphological metrics. While there have been multiple robust studies comparing different imaging metrics (Hutton et al., 2009; Im et al., 2008; Lövdén et al., 2013; Pantazis et al., 2010; Shimony et al., 2016; Wang et al., 2019; Wierenga et al., 2014), few have included the breadth of morphometry assessed here. Our approach, therefore, allowed us to directly compare the magnitude of cortical age-related differences and changes across a range of metrics.

The biggest age-related change (cross-sectionally and longitudinally) was that of cortical thickness, followed (cross-sectionally) by curvature. This suggests that the most striking structural transformation the human brain undergoes with age – at least of those detectable with MRI – is that the cortex thins while also becoming more ‘curved’. The width and depth of cortical sulci might influence the complexity metric, such that more atrophied brains might exhibit an increase in gyral complexity but not a decrease in surface area (Narr, et al., 2004; Lemaitre et al., 2012).

We also show that *combining* shape measures outperforms any individual metrics’ ability to capture age-related and cognitive differences: together, the eight morphometric metrics assessed here explained almost double the variance compared to that captured by thickness and surface area alone. Thus, the fact that multiple morphometric measures provided partially complementary information about the outcome highlights the potential usefulness in assessing various morphological shape measures when investigating the ageing brain and cognitive abilities.

### Methodological strengths and limitations

In addition to the large sample size and the assessment of multiple shape metrics, the integration of cross-sectional and longitudinal data is of note. Recent reviews and commentaries have pointed to the limitations of cross-sectional analyses when investigating brain-cognition relationships in the ageing brain (see Oschwald 2020 for a discussion). While we agree that collecting longitudinal data is almost always preferable, we acknowledge that it is not always attainable. Our approach of integrating cross-sectional and longitudinal data, where the latter largely confirmed the findings of the former, offers some validation of cross-sectional approaches.

Another key strength of this paper is that we successfully replicated our cross-sectional and longitudinal findings in an independent cohort. In doing so, we not only validated the apparent existence of the morphological double dissociation, but showed that it is not subject to specific features of the Cam-CAN data. Indeed, replicating our results despite important differences between the two datasets increases the robustness of our findings considerably. For instance, the cognitive tests differed (Cattell in Cam-CAN, WASI Matrix in LCBC), suggesting that surface area captures the broader construct of fluid intelligence (rather than test-specific features). Moreover, while the morphological metrics assessed in our initial Cam-CAN study offered an intriguing description of the ageing brain, obtaining them required five separate processing pipelines (FreeSurfer (Fischl, 2012), FreeSurfer Long (Reuter et al., 2012), Mindboggle (Klein et al., 2017), SPM (Ashburner & Friston, 2000) and the Fractal Dimensionality Toolbox calcFD (Madan & Kensinger, 2016)). The fact that our results replicated in canonical metrics (all of which are part of the standard FreeSurfer output) might lower the threshold for future research to, where appropriate, investigate surface area and cortical thickness separately.

The breadth of structural brain metrics reviewed in this paper also comes with some important limitations. First, we were not able investigate the *changes* of several of the metrics which we had assessed in our cross-sectional analyses. This is because the pipelines used to calculate these additional metrics (e.g. Mindboggle) are not yet optimised for longitudinal data. Particularly curvature, which showed a very strong age effect cross-sectionally, would have been interesting to explore longitudinally. Likewise, fractal dimensionality, which measures cortical complexity and correlated strongly with age *and* cognition in our cross-sectional analyses, might be a promising candidate for future longitudinal investigations.

## Conclusion

In this paper, we found cross-sectional and longitudinal evidence for a brain-cognition double dissociation: two morphological metrics, surface area and cortical thickness, which tend to be investigated together through grey matter volume, are differentially associated with age and fluid intelligence: while thickness is strongly associated with age, it has weak associations with change in fluid intelligence – a pattern that is reversed for surface area, which captures cognitive change and difference well, and age relatively poorly. We therefore recommend that rather than using grey matter volume as the default measure, researchers should choose structural brain metrics depending on the question under investigation. Doing so will allow us to advance our understanding of the functional significance of these dissociable aspects of brain morphology.

## Methods

### Initial Cohort

#### Participants

Participants were drawn from the Cambridge Centre for Ageing and Neuroscience (Cam-CAN) study, which has been described in more detail elsewhere (Shafto et al., 2014; Taylor et al., 2017). 708 healthy adults (359 women, 349 men) from the larger cohort were scanned, with approximately 100 people in each decade (age range 18-88, Mean=53.4, Standard Deviation (sd) = 18.62). We used calendar age (years) as a measure of participants’ age. Cognitive ability was measured using the Cattell Culture Fair test of fluid intelligence (Cattell, 1971). For an age-independent measure of cognition, we calculated age-residualized fluid intelligence scores by regressing the Cattell raw scores on age (see Borgeest et al., 2019). Residuals adjust for age-expected declines, allowing, for example, an 80-year-old person with a relatively low absolute score to be considered cognitively healthier than a younger individual with a higher score.

A subset of participants (N=261) was scanned twice, with an average interval between the first and the second scan of 1.33 years (sd = 0.66). Additionally, a (partially separate) subset of participants (N=233) completed the Cattell test twice with an average interval between the two cognitive tests of 6.0 years (sd = 0.67). Two waves of both brain *and* cognitive data were available for 115 participants.

#### Imaging data acquisition and pre-processing

T1- and T2-weighted 1 mm isotropic magnetic resonance imaging scans were available for 647 participants (Taylor et al., 2017). To ensure the quality of the image segmentations, we adapted a recently developed supervised learning tool (Klapwijk et al., 2019), which led us to exclude six participants due to low-quality segmentations. Our quality control process is described further the supplementary materials. In order to investigate (cross-sectional) brain morphology in as much detail as possible, we examined a total of eight brain metrics: in addition to three FreeSurfer-derived measures of cortical volume, thickness and surface area (derived from a standard FreeSurfer recon-all pipeline), we examined grey-matter volume derived from SPM 12 (voxel-based morphometry which includes sub-cortical grey-matter too, while FreeSurfer includes only cortical estimates) and four additional morphological measures: from *Mindboggle* (see Klein et al. 2017 for more detail) we derived sulcal depth, curvature and “thickinthehead” (a recently developed cortical thickness measure that avoids FreeSurfer’s reconstruction-based limitations); and from the *calcFD* toolbox (Madan & Kensinger, 2016) we calculated fractal dimensionality as a measure of cortical complexity. To extract reliable brain structure estimates from the longitudinal subsample, images were automatically processed with FreeSurfer’s longitudinal stream (Reuter et al., 2012). This yielded co-registered measures of volume, cortical thickness and surface area for the two waves. Note that we did not explore the other morphological metrics longitudinally because the *Mindboggle* and *calcFD* pipeline are not currently optimised for longitudinal data (see discussion). Brain regions were defined according to the Desikan-Killiany-Tourville (DKT) protocol, which yields 62 brain regions (Klein & Tourville, 2012).

#### Cross-sectional analyses

All analyses were carried out using R (R Core Team, 2013), and the code used for this paper is available on the Open Science Framework (https://osf.io/n6b4j/).

First, we calculated whole brain as well as regional correlations between each metric and age, fluid intelligence and age-residualized fluid intelligence. Regional correlations were FDR corrected at alpha = 0.05. Next, we estimated a series of path models to assess which combination of whole brain metrics best predicted age, fluid intelligence and age-residualized fluid intelligence. We then examined the robustness of our frequentist modelling approach with a Bayesian modelling framework (see supplementary materials).

#### Longitudinal analyses

To assess neural and fluid intelligence change between time point 1 and time point 2, we fit a series of longitudinal structural equation models for each longitudinal FreeSurfer metric (whole brain volume, thickness and surface area) and fluid intelligence. Before assessing cognitive change, we also tested for longitudinal measurement invariance (Widaman et al., 2010). Additionally, as the second Cattell test was completed online by approximately half of the participants, versus pencil and paper by the other half, we investigated whether these two groups differed in their measurement properties by assessing metric invariance (constraining factor loadings) and scalar invariance (constraining intercepts).

To understand whether cognitive change was correlated with morphometric change, and if so, whether this relationship differed for the different cortical metrics, we extracted and estimated the rates of cognitive and brain structure change in a series of second order latent change score models (Ferrer et al., 2008; Ferrer & McArdle, 2010; McArdle & Hamagami, 2001; McArdle & Nesselroade, 2003). Second order latent change score models (2LCSM) first estimate latent factors at each time point, and then estimate latent change over time. Steiger’s Z-Tests were performed to assess whether the change-change relationships differed significantly between the different metrics (Steiger, 1980). Given that properties of the data, obtaining latent cognitive scores was not possible in the replication sample (see below), so we also ran the models with observed variables only within Cam-CAN to ensure maximal comparability between the two sets of analyses. We ran models on participants with at least one cognitive score (N=362) using full information maximum likelihood (FIML, which assumes data are missing-at-random, Enders & Mansolf, 2018, and enables robust standard errors to account for missingness).

### Replication Cohort

To assess the robustness of our results, we investigated whether our core findings replicated in a second, independent dataset. To this end, we analysed data from the Centre for Lifespan Changes in Brain and Cognition at the University of Oslo (LCBC; https://www.oslobrains.no/), which is part of the European Lifebrain project (Walhovd et al., 2018) together with Cam-CAN and other publicly available datasets. The LCBC data consist of a collection of studies, which have been described elsewhere (Walhovd et al., 2016). Briefly, our analyses included 1236 adults aged 18-93 years (median = 37, sd = 20.64). We used WASI Matrix (raw scores) as our measure of fluid intelligence because it is most similar to the Cattell task assessed in Cam-CAN. FreeSurfer-derived cortical thickness, volume and surface area served as our morphological measures (for details on cross-sectional and longitudinal image acquisition and pre-processing see (Walhovd et al., 2016)). At least two waves of cognitive and/or neural data were available for 389 participants. Where participants had more than two waves, we selected their first and last time point, maximizing the interval between waves as well as the data similarity between samples. This allowed us to include the largest possible number of participants in our longitudinal analyses while maintaining two-wave models comparable to those described in Cam-CAN. The mean interval between the two waves so defined was 5.18 years (min = 0.73, max = 10.0, sd = 2.59 years).

Our analysis pipeline mirrored that described above: cross-sectionally, whole brain correlations were followed by frequentist path models and Bayesian model selection analyses. Longitudinally, LCSMs assessed cognitive and neural change separately; and we ran a series 2LCSMs to investigate the relationship between cognitive change and neural change. The FIML models included 722 participants. Note that it was not possible to derive latent cognitive factor scores for the longitudinal models as individual WASI scores were not available, so the LCBC longitudinal models used observed cognitive variables (but were otherwise identical to Cam-CAN models). The LCSM data and analyses are described in more detail in the supplementary material.

## Borgeest et al. (2021): Supplementary Materials

### 1. Descriptive Statistics

**Table 1:** Descriptive statistics for Cam-CAN data

| Cam-CAN | N | Mean | SD | Median | Min | Max | Skew | Kurtosis |
| --- | --- | --- | --- | --- | --- | --- | --- | --- |
| Age | 641 | 54.04 | 18.56 | 54.00 | 18.00 | 88.00 | -0.05 | -1.15 |
| WB Volume | 641 | 7114.64 | 899.87 | 7034.19 | 930.07 | 4939.80 | 0.36 | -0.17 |
| WB Area | 641 | 3177.87 | 320.97 | 3157.00 | 2442.25 | 4445.75 | 0.38 | 0.09 |
| WB Thickness | 641 | 2.66 | 0.12 | 2.67 | 2.19 | 2.98 | -0.50 | 0.87 |
| Cattell | 622 | 31.05 | 6.74 | 33.00 | 11.00 | 44.00 | -0.56 | -0.16 |

**Table 2:** Descriptive statistics for LCBC data

| LCBC | N | Mean | SD | Median | Min | Max | Skew | Kurtosis |
| --- | --- | --- | --- | --- | --- | --- | --- | --- |
| Age | 1236 | 41.55 | 20.32 | 31.95 | 18.0 | 93.35 | 0.71 | -1.02 |
| WB Volume | 1188 | 7453.09 | 853.41 | 7441.81 | 890.39 | 5092.91 | 0.14 | -0.41 |
| WB Area | 1188 | 2630.76 | 246.85 | 2618.33 | 1859.63 | 3300.62 | 0.15 | -0.32 |
| WB Thickness | 1199 | 2.60 | 0.11 | 2.61 | 2.09 | 2.91 | -0.38 | -0.03 |
| Wasi Matrix Raw | 1234 | 27.67 | 4.64 | 20.00 | 6.00 | 35.00 | -.69 | 4.06 |

**Figure 1:**
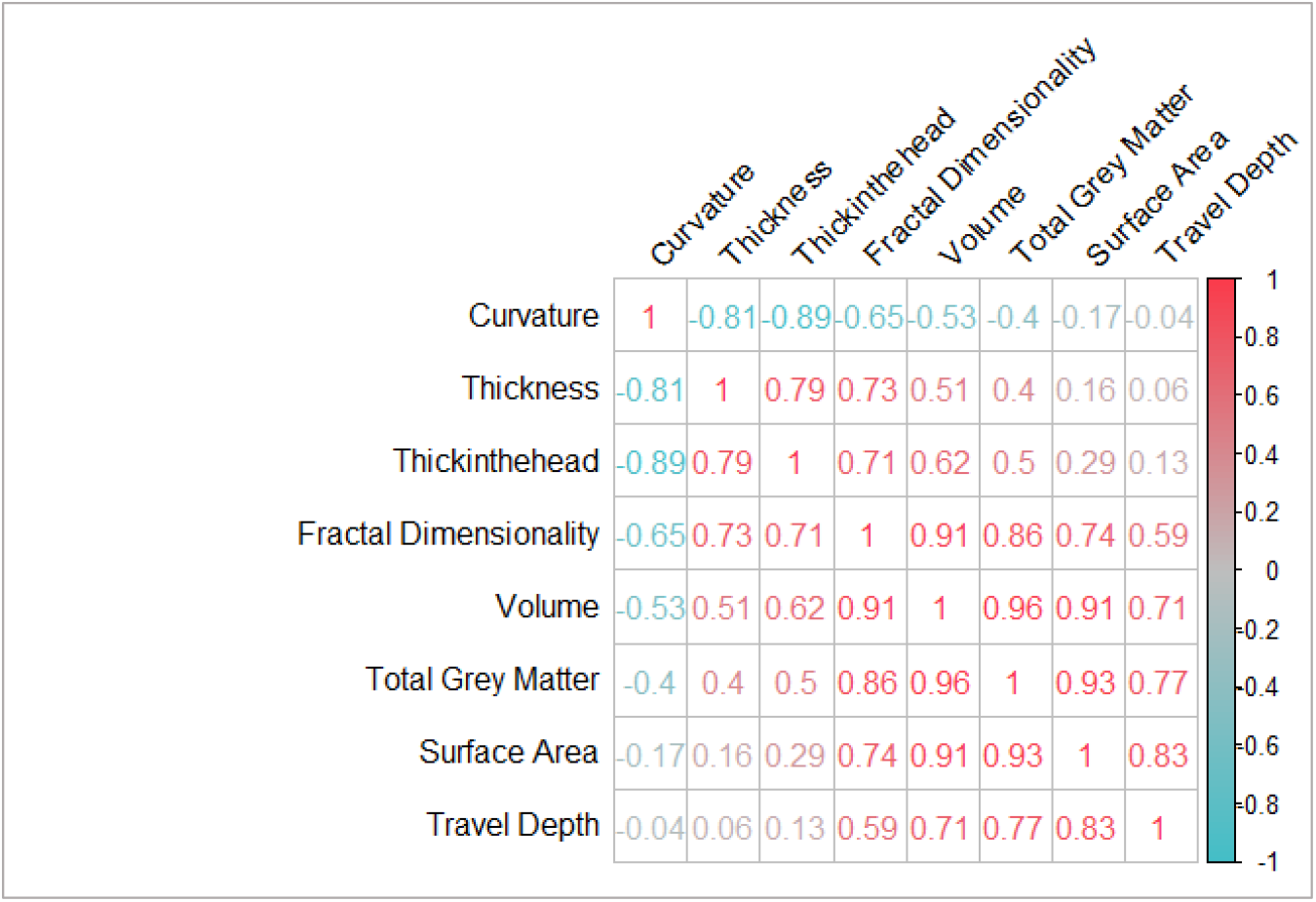
Correlation matrix of the eight brain structure metrics. Note that surface area and thickness are correlated r = 0.16

**Figure 2:**
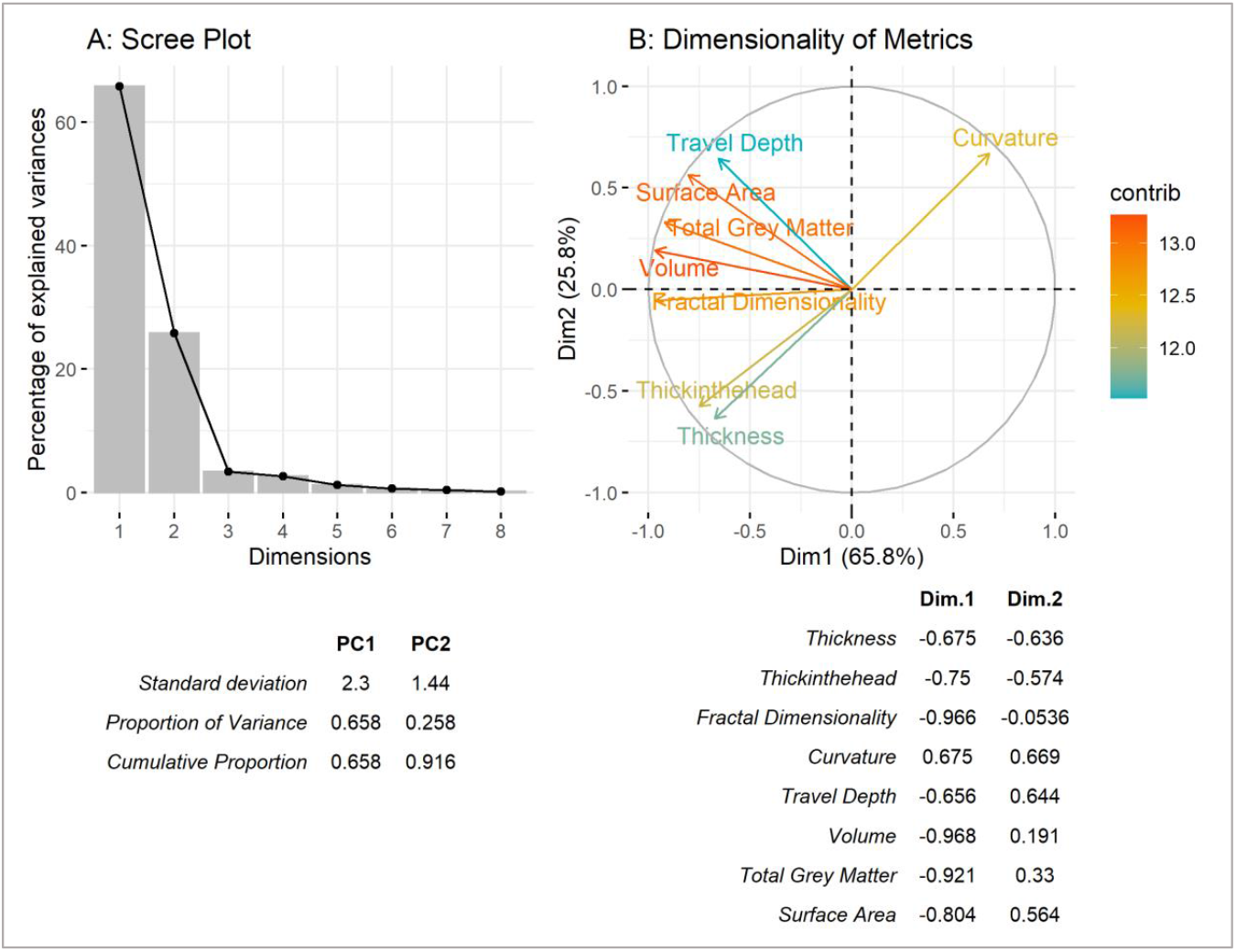
Results of Principal Component Analysis

### 2. Imaging data acquisition and pre-processing

We based our quality control process on the supervised learning tool ‘Qoala-T’ developed by Klapwijk et al., which was originally developed for child and adolescent samples (see manual, 2019a, and manuscript, 2019b). First, we manually rated the quality of 12% of our FreeSurfer preprocessed Cam-CAN scans, thereby surpassing the proportion of 10% as recommended by the Qoala-T authors. These scans later served as input for Qoala-T, so the algorithm would learn to distinguish between scan qualities suitable or unsuitable for further analyses. Second, following the manual ratings, we used Qoala-T’s publicly available quality control tool to assess the quality of all T1 CamCAN images. This resulted in six participants being excluded from the sample (age 32 – 71, median = 59).

We have uploaded a detailed rating procedure to this project’s OSF page **<u>(link here)</u>** as we hope that it will help other researchers implement versions of this semi-automatic quality control procedure for large adult lifespan samples.

### 2 Whole Brain Correlations

**Table 3:**
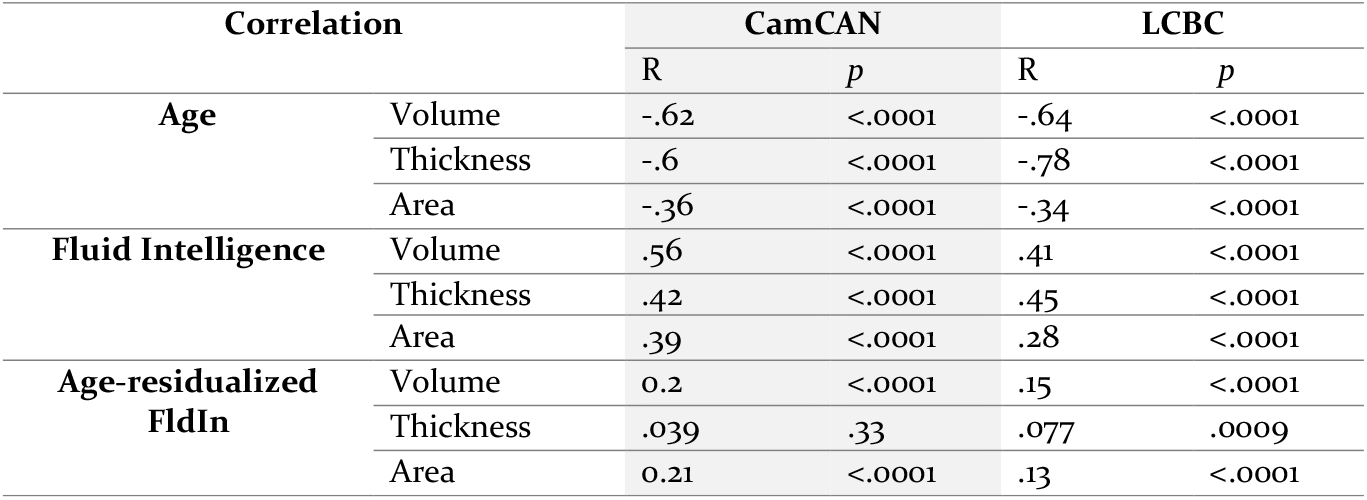
Comparing whole brain correlations in CamCAN and LCBC data

**Table 4:** Comparing linear and quadratic model fit for the metric-age correlations in CamCAN. The best fitting model (with lower BIC) is marked with *.

| Metric | Model | R-Squared | F-Statistic | p | BIC |
| --- | --- | --- | --- | --- | --- |
| Cortical Volume | Linear | 0.38 | 399.7 | <0.001 | 5270.861 |
|  | Quadratic * | 0.39 | 206.2 | <0.001 | 5269.154 |
| Cortical Thickness | Linear * | 0.36 | 366.2 | <0.001 | 5291.885 |
|  | Quadratic | 0.37 | 184.2 | <0.001 | 5296.544 |
| Surface Area | Linear * | 0.13 | 96.94 | <0.001 | 5491.733 |
|  | Quadratic | 0.13 | 48.56 | <0.001 | 5497.909 |
| Thickinthehead | Linear * | 0.71 | 1538 | <0.001 | 4796.678 |
|  | Quadratic | 0.71 | 768.2 | <0.001 | 4802.762 |
| Curvature | Linear | 0.60 | 955.2 | <0.001 | 4996.165 |
|  | Quadratic * | 0.63 | 532.4 | <0.001 | 4959.439 |
| Sulcal Depth | Linear * | 0.14 | 106.2 | <0.001 | 5483.685 |
|  | Quadratic | 0.14 | 53.17 | <0.001 | 5489.911 |
| Grey Matter Volume (SPM) | Linear * | 0.30 | 269.4 | <0.001 | 5356.77 |
|  | Quadratic | 0.30 | 135.2 | <0.001 | 5361.933 |
| Fractal Dimensionality | Linear * | 0.42 | 467.6 | <0.001 | 5230.34 |
|  | Quadratic | 0.42 | 234.1 | <0.001 | 5235.915 |

### 3 Frequentist modelling approach

We examined whether the different metrics of brain structure provided unique and complementary information about age and cognitive ability. To do so, we ran frequentist path models and Bayesian model selection framework in which cortical thickness and surface area predicted either age, fluid intelligence or age-adjusted fluid intelligence (ignoring volume since this is the product of thickness and surface area). These revealed that the best model of age and fluid intelligence required both surface area and thickness (Figure 1 A-B). In contrast, individual differences in (age-residualized) fluid intelligence were best captured by surface area alone (Figure 1 C). These models explained 44, 29 and 4 percent of the variance of age, fluid intelligence and age-residualized fluid intelligence, respectively. The Bayesian model selection – which led to identical conclusions – is plotted in the supplementary materials.

The full models that included all 8 metrics are depicted in Figure 1 D-F. The total variance explained by these models was 76, 46 and 7 percent for age, fluid intelligence and age-residualized fluid intelligence, respectively – almost double the variance explained by thickness and area alone. Moreover, the fact that multiple morphometric measures provided partially complementary information about the outcome highlights the potential usefulness in assessing various morphological shape measures when investigating the ageing brain and cognitive abilities. As was the case for the first set of models, the Bayesian model selection arrived at the same conclusions as the frequentist model selection (see supplementary materials): For age, the best model included Thickness, Thickinthehead, Curvature, TGM and Surface Area. Fluid intelligence was best captured by Thickinthehead, Curvature, TGM, Surface Area, Thickness and FD. Finally, the best model for age-residualized fluid intelligence included Fractal Dimensionality and Thickness. Interestingly, when FD was not included in the models, the best model for age-residualized fluid intelligence included surface area only, suggesting that surface area and FD capture similar variance.

**Figure 3:**
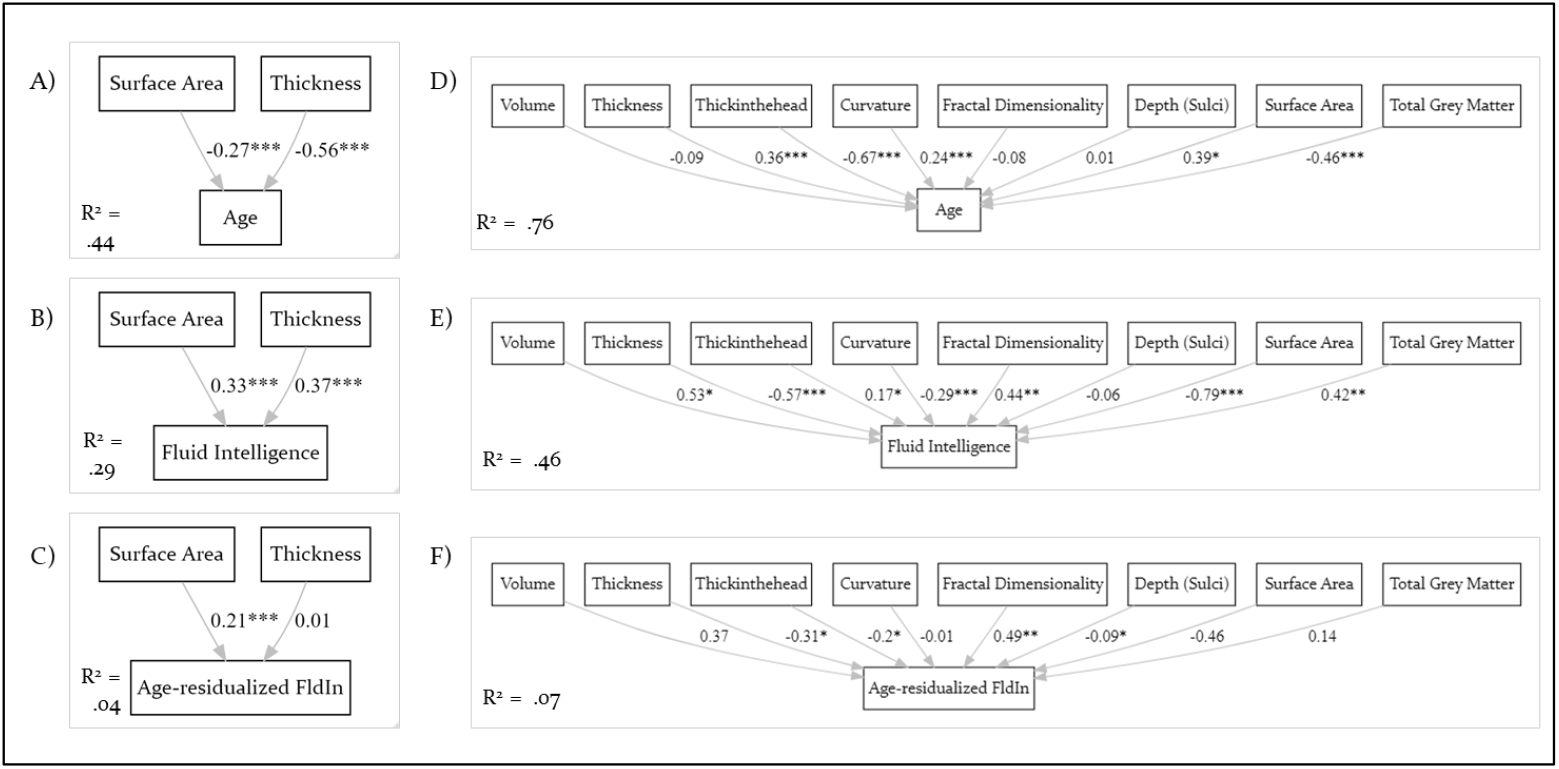
CamCAN path model results. Models A-C were for the area and thickness only, models D-F included all eight brain structure metrics.

### 4 Bayesian model selection

We validated our frequentist modelling approach with a Bayesian modelling framework (Rouder et al., 2012) using Bayesian regression. As before in this cohort (Gadie et al., 2017), we used the default, symmetric Cauchy prior with width of 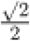 which translates to a 50% confidence that the true effect will lie between −0.707 and 0.707. Doing so yields a Bayes factor for all possible subsets of predictors, thus yielding the model that optimally balances parsimony (excluding unnecessary predictors) with prediction power.

All Bayesian models confirmed the frequentist ones. For age, the best model was comprised of Thickinthehead, Curvature, TGM, Surface Area and Thickness (Figure 2). Fluid intelligence was best captured by Curvature, TGM, Surface Area, Thickness, FD and Volume (Figure 3). Finally, age-residualized fluid intelligence was best predicted by FD and Thickness (Figure 4).

**Figure 4:**
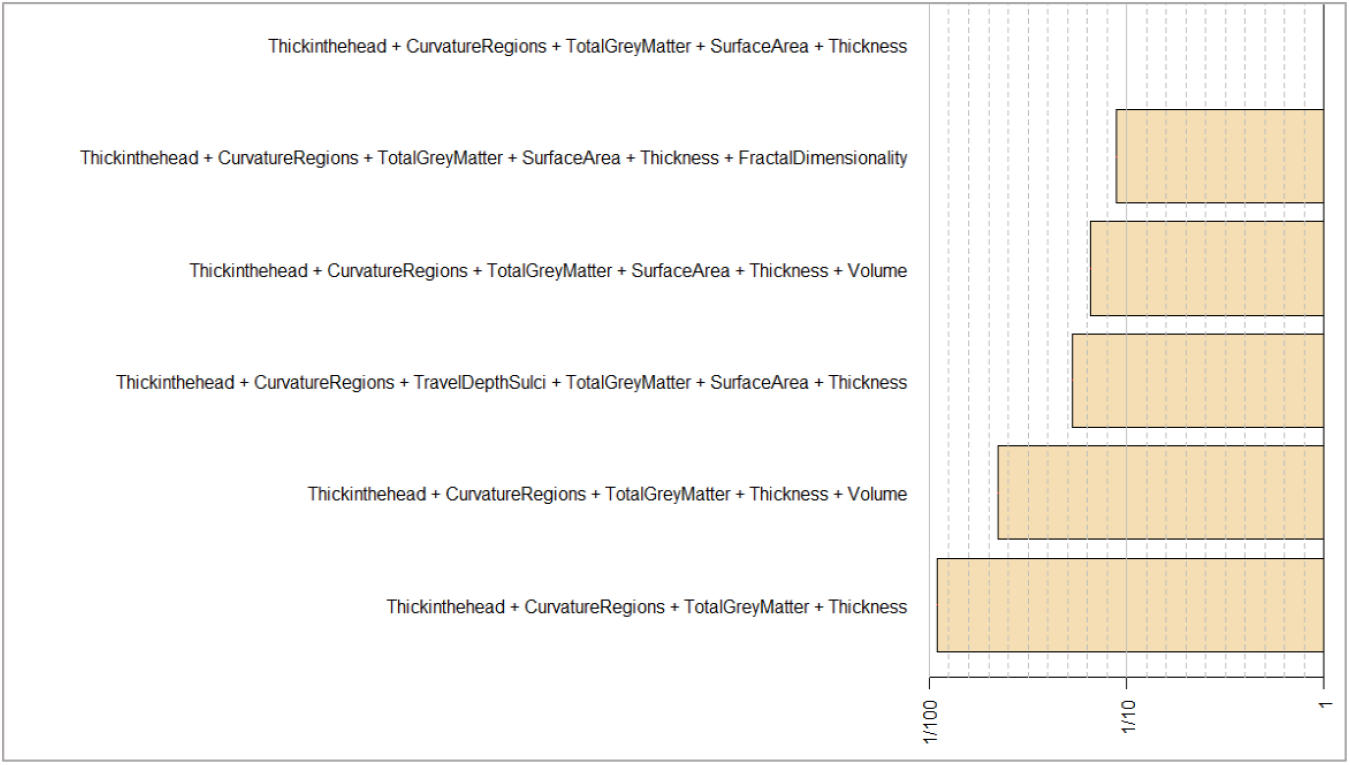
Bayesian model selection framework, predicting Age in CamCAN. Compares the best model (top row) to the next five best fitting models.

**Figure 5:**
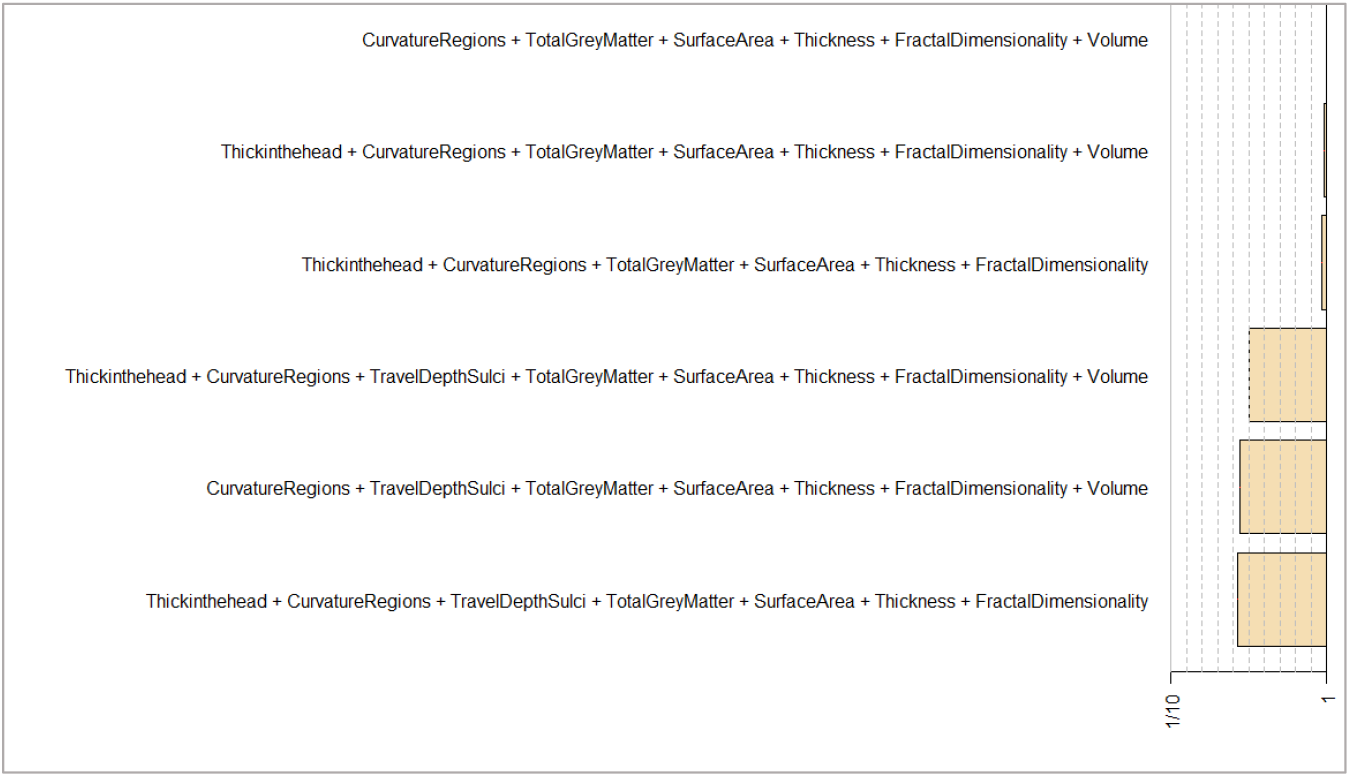
Bayesian model selection framework, predicting fluid intelligence in CamCAN. Compares the best model (top row) to the next five best fitting models.

**Figure 6:**
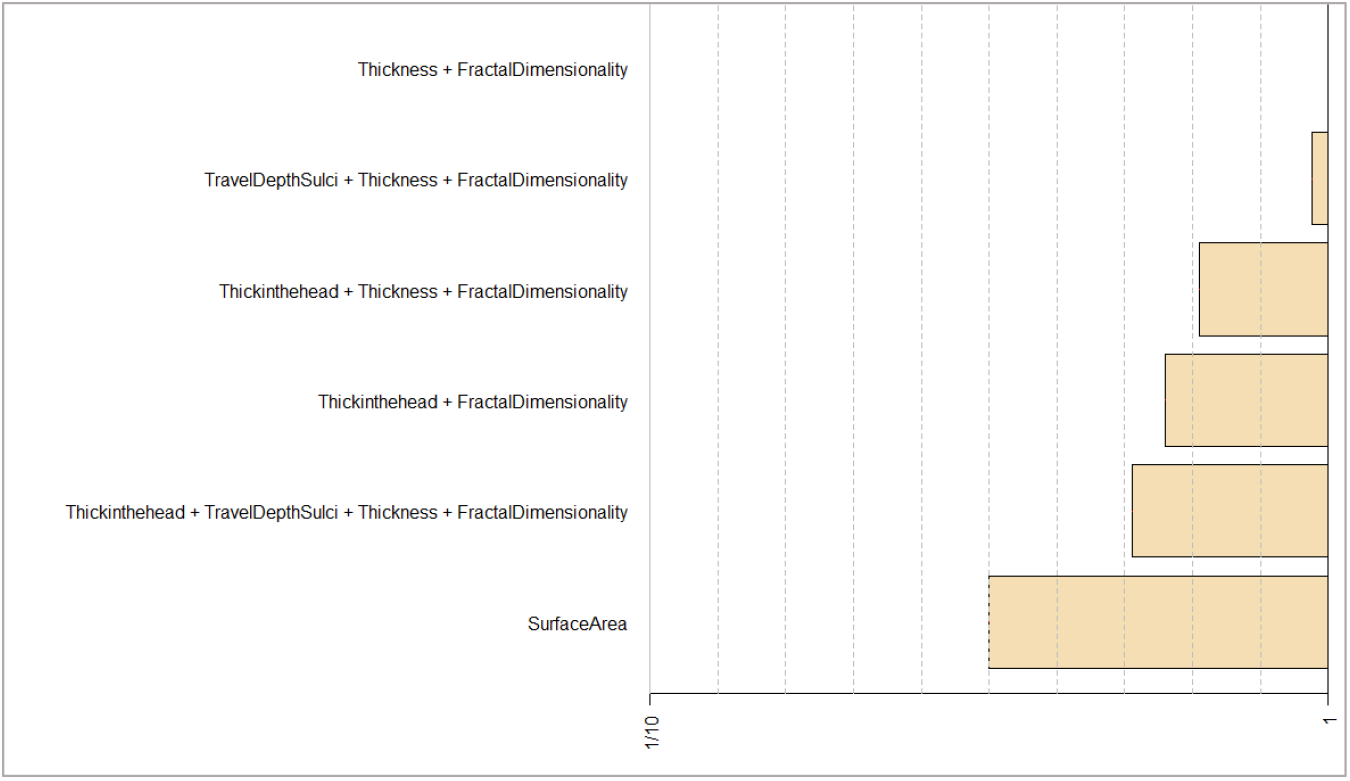
Bayesian model selection framework, predicting age-residualized fluid intelligence in CamCAN. Compares the best model (top row) to the next five best fitting models.

### 5 Regional results

In Cam-CAN, after looking at whole brain correlations between the eight metrics and age, fluid intelligence and age-residualized fluid intelligence, we investigated regional correlations. Regions were assigned 62 labels following the Desikan-Killiany-Tourville (DKT) protocol in the Mindboggle pipeline (Klein et al., 2018). We then averaged across both hemispheres. Results are shown in Tables 4–6 and plotted in Figures 4–6. Note that data for the entorhinal, banks superior temporal and temporal pole was only available for Thickinthehead and Volume.

Our regional investigations further supported the morphological dichotomy found in the whole brain analyses. For cortical thickness, all 32 brain regions (averaged across the hemispheres) were significantly correlated with age (all correlations were FDR corrected at alpha = 0.05), while not a single region predicted age-residualized fluid intelligence (Figure 3 and Tables 4–6 in supplementary materials). In contrast, for surface area, *all* regions were significantly associated with age-residualized fluid intelligence. While regional surface area also correlated with age, the correlations were substantially weaker than the brain-age correlations for cortical thickness.

The precentral gyrus was the region with the strongest age effects in five out of eight metrics: curvature (r=.74), thickness (r=-.66), thickinthehead (r=-.87), volume (r=-.71), TGM (r=.-66). More regional results are shown in Tables 4–6 and Figures 4–6 in the supplementary materials.

**Figure 7:**
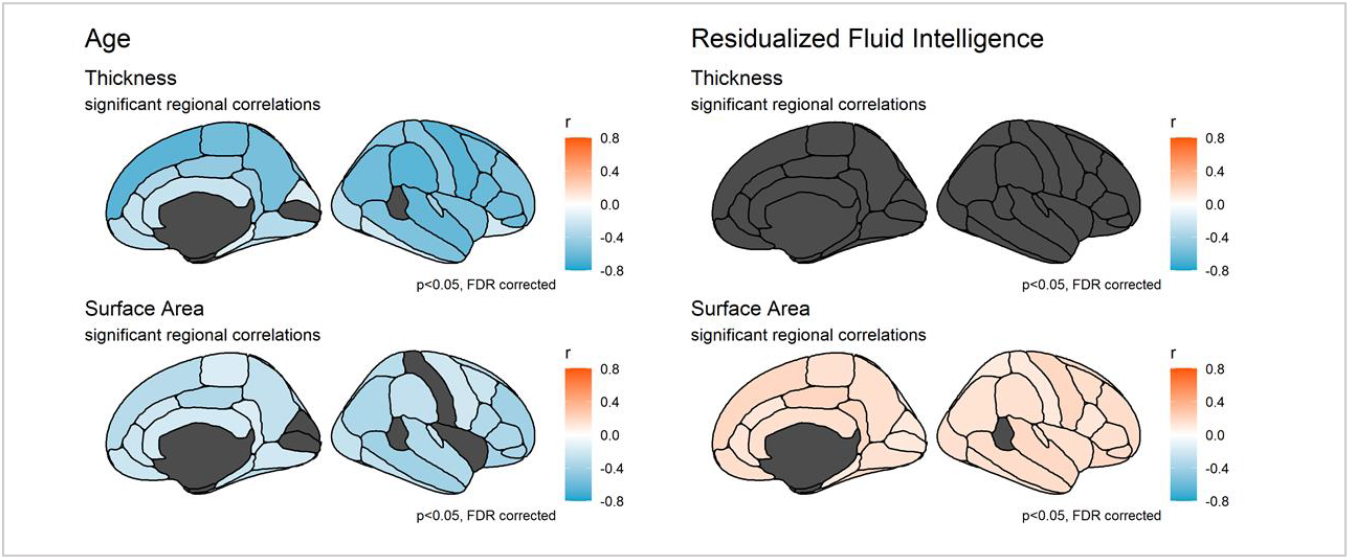
Significant regional age- and age-residualized fluid intelligence correlations. Correlations are FDR corrected at alpha = 0.05. Shows a double dissociation, whereby cortical thickness predicts age and not cognition, and vice versa for surface area. Note that grey indicates non-significant or missing regions.

**Figure 8:**
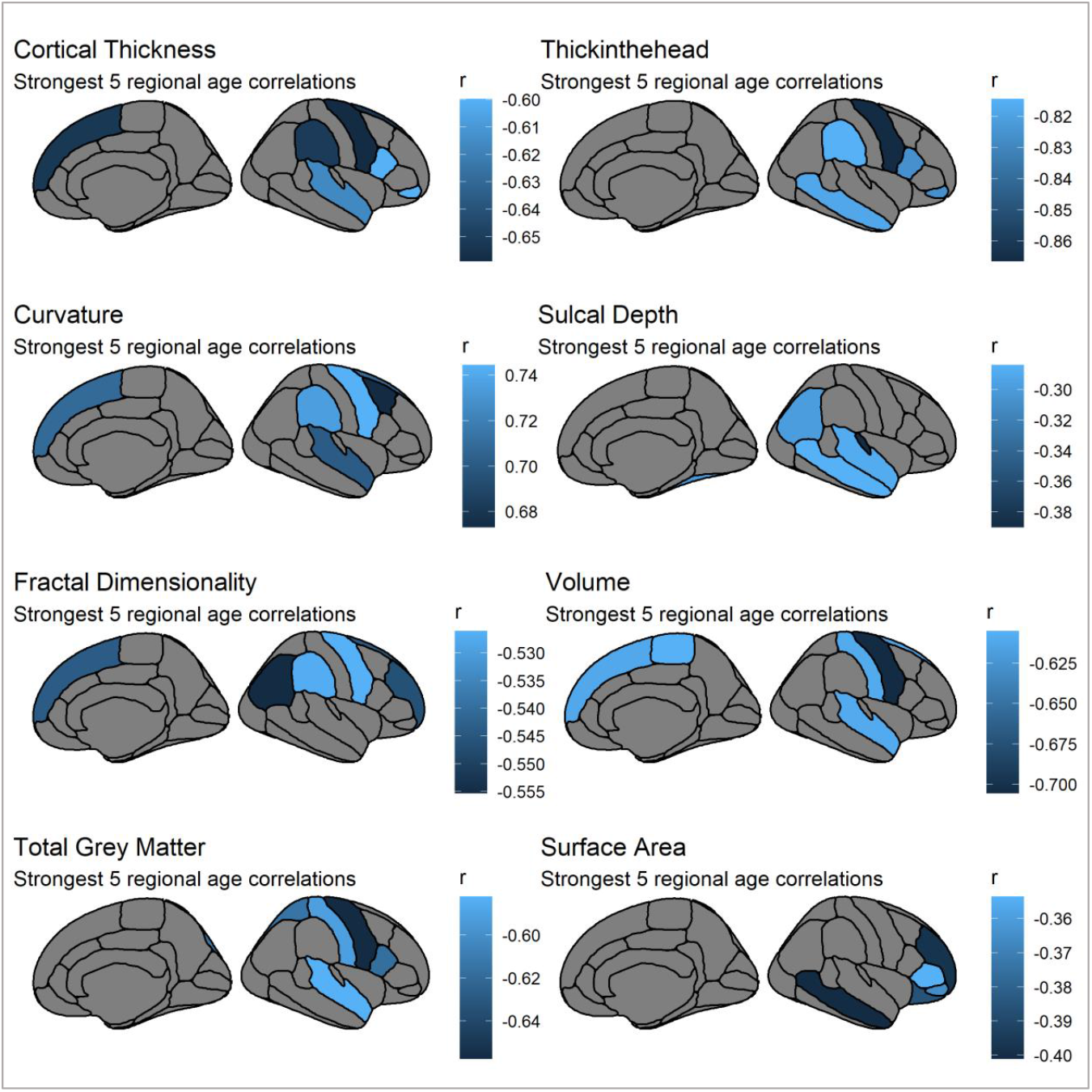
regions most strongly associated with age. Shows a large variability, with volume showing pre-frontal age effects while, for instance, sulcal depth effects are focused in the temporal lobes.

**Table 5:** Regional age correlations in Cam-CAN. All p-values are FDR corrected at alpha = 0.05.

|  | Fractal Dim. |  | Curvature |  | Thickness |  | Thickinthhead |  | Volume |  | TGM |  | Depth |  | Area |  |
| --- | --- | --- | --- | --- | --- | --- | --- | --- | --- | --- | --- | --- | --- | --- | --- | --- |
| ROI | r | p | r | p | r | p | r | p | r | p | r | p | r | p | r | p |
| bankssts | NA | NA | NA | NA | NA | NA | -0.783 | <.001 | -0.496 | <.001 | NA | NA | NA | NA | NA | NA |
| caudal anterior cingulate | -0.285 | <.001 | 0.589 | <.001 | -0.353 | <.001 | -0.649 | <.001 | -0.366 | <.001 | -0.443 | <.001 | 0.131 | 0.002 | -0.216 | <.001 |
| caudal middle frontal | -0.479 | <.001 | 0.673 | <.001 | -0.569 | <.001 | -0.772 | <.001 | -0.501 | <.001 | -0.557 | <.001 | -0.118 | 0.005 | -0.233 | <.001 |
| corpus callosum | -0.207 | <.001 | 0.451 | <.001 | -0.239 | <.001 | -0.609 | <.001 | -0.361 | <.001 | -0.536 | <.001 | -0.033 | 0.494 | -0.191 | <.001 |
| cuneus | -0.037 | 0.349 | -0.036 | 0.365 | -0.159 | <.001 | -0.357 | <.001 | 0.016 | 0.685 | -0.35 | <.001 | -0.008 | 0.894 | -0.009 | 0.843 |
| entorhinal | NA | NA | NA | NA | NA | NA | -0.313 | <.001 | -0.264 | <.001 | NA | NA | NA | NA | NA | NA |
| fusiform | -0.38 | <.001 | 0.492 | <.001 | -0.305 | <.001 | -0.683 | <.001 | -0.374 | <.001 | -0.461 | <.001 | -0.298 | <.001 | -0.306 | <.001 |
| inferior parietal | -0.555 | <.001 | 0.671 | <.001 | -0.585 | <.001 | -0.747 | <.001 | -0.558 | <.001 | -0.524 | <.001 | -0.298 | <.001 | -0.347 | <.001 |
| inferior temporal | -0.27 | <.001 | 0.475 | <.001 | -0.209 | <.001 | -0.646 | <.001 | -0.293 | <.001 | -0.431 | <.001 | -0.216 | <.001 | -0.268 | <.001 |
| insula | -0.242 | <.001 | 0.63 | <.001 | -0.536 | <.001 | -0.71 | <.001 | -0.423 | <.001 | -0.49 | <.001 | 0.018 | 0.769 | -0.004 | 0.929 |
| isthmus cingulate | -0.352 | <.001 | 0.523 | <.001 | -0.423 | <.001 | -0.76 | <.001 | -0.405 | <.001 | -0.387 | <.001 | -0.048 | 0.303 | -0.165 | <.001 |
| lateral occipital | -0.414 | <.001 | 0.519 | <.001 | -0.297 | <.001 | -0.647 | <.001 | -0.329 | <.001 | -0.467 | <.001 | -0.176 | <.001 | -0.254 | <.001 |
| lateral orbitofrontal | -0.389 | <.001 | 0.396 | <.001 | -0.2 | <.001 | -0.627 | <.001 | -0.491 | <.001 | -0.502 | <.001 | -0.226 | <.001 | -0.386 | <.001 |
| lingual | -0.256 | <.001 | 0.506 | <.001 | -0.321 | <.001 | -0.65 | <.001 | -0.343 | <.001 | -0.567 | <.001 | -0.14 | 0.001 | -0.201 | <.001 |
| medial orbitofrontal | -0.239 | <.001 | 0.335 | <.001 | -0.296 | <.001 | -0.55 | <.001 | -0.443 | <.001 | -0.541 | <.001 | -0.002 | 0.972 | -0.226 | <.001 |
| middle temporal | -0.446 | <.001 | 0.637 | <.001 | -0.534 | <.001 | -0.818 | <.001 | -0.544 | <.001 | -0.513 | <.001 | -0.284 | <.001 | -0.401 | <.001 |
| paracentral | -0.463 | <.001 | 0.459 | <.001 | -0.578 | <.001 | -0.663 | <.001 | -0.605 | <.001 | -0.564 | <.001 | -0.039 | 0.409 | -0.161 | <.001 |
| parahippocampal | -0.116 | 0.003 | 0.226 | <.001 | -0.149 | <.001 | -0.45 | <.001 | -0.354 | <.001 | -0.433 | <.001 | -0.062 | 0.17 | -0.232 | <.001 |
| pars opercularis | -0.487 | <.001 | 0.637 | <.001 | -0.6 | <.001 | -0.826 | <.001 | -0.597 | <.001 | -0.617 | <.001 | -0.063 | 0.17 | -0.333 | <.001 |
| pars orbitalis | -0.397 | <.001 | 0.252 | <.001 | -0.6 | <.001 | -0.826 | <.001 | -0.459 | <.001 | -0.523 | <.001 | -0.077 | 0.085 | -0.365 | <.001 |
| pars triangularis | -0.508 | <.001 | 0.564 | <.001 | -0.581 | <.001 | -0.797 | <.001 | -0.599 | <.001 | -0.525 | <.001 | -0.124 | 0.003 | -0.354 | <.001 |
| pericalcarine | -0.118 | 0.003 | 0.487 | <.001 | -0.049 | 0.213 | -0.604 | <.001 | -0.389 | <.001 | -0.46 | <.001 | -0.015 | 0.775 | -0.055 | 0.178 |
| postcentral | -0.469 | <.001 | 0.632 | <.001 | -0.494 | <.001 | -0.773 | <.001 | -0.609 | <.001 | -0.591 | <.001 | -0.218 | <.001 | -0.055 | 0.178 |
| posterior cingulate | -0.45 | <.001 | 0.595 | <.001 | -0.459 | <.001 | -0.706 | <.001 | -0.529 | <.001 | -0.522 | <.001 | 0.169 | <.001 | -0.341 | <.001 |
| precentral | -0.526 | <.001 | 0.744 | <.001 | -0.659 | <.001 | -0.867 | <.001 | -0.706 | <.001 | -0.658 | <.001 | -0.105 | 0.014 | -0.205 | <.001 |
| precuneus | -0.487 | <.001 | 0.663 | <.001 | -0.559 | <.001 | -0.731 | <.001 | -0.526 | <.001 | -0.408 | <.001 | 0.052 | 0.259 | -0.266 | <.001 |
| rostral anterior cingulate | -0.316 | <.001 | 0.379 | <.001 | -0.248 | <.001 | -0.597 | <.001 | -0.36 | <.001 | -0.53 | <.001 | -0.017 | 0.769 | -0.227 | <.001 |
| rostral middle frontal | -0.546 | <.001 | 0.597 | <.001 | -0.512 | <.001 | -0.674 | <.001 | -0.583 | <.001 | -0.56 | <.001 | -0.265 | <.001 | -0.398 | <.001 |
| superior frontal | -0.544 | <.001 | 0.709 | <.001 | -0.653 | <.001 | -0.759 | <.001 | -0.611 | <.001 | -0.523 | <.001 | 0.001 | 0.972 | -0.313 | <.001 |
| superior parietal | -0.514 | <.001 | 0.595 | <.001 | -0.491 | <.001 | -0.62 | <.001 | -0.562 | <.001 | -0.614 | <.001 | -0.071 | 0.114 | -0.298 | <.001 |
| superior temporal | -0.446 | <.001 | 0.701 | <.001 | -0.616 | <.001 | -0.62 | <.001 | -0.609 | <.001 | -0.582 | <.001 | -0.288 | <.001 | -0.332 | <.001 |
| supramarginal | -0.527 | <.001 | 0.735 | <.001 | -0.651 | <.001 | -0.814 | <.001 | -0.532 | <.001 | -0.529 | <.001 | -0.131 | 0.002 | -0.266 | <.001 |
| temporal pole | NA | NA | NA | NA | NA | NA | -0.469 | <.001 | -0.065 | 0.1 | NA | NA | NA | NA | NA | NA |
| transverse temporal | -0.441 | <.001 | 0.554 | <.001 | -0.403 | <.001 | -0.772 | <.001 | -0.523 | <.001 | -0.555 | <.001 | -0.39 | <.001 | -0.251 | <.001 |

**Table 6:**
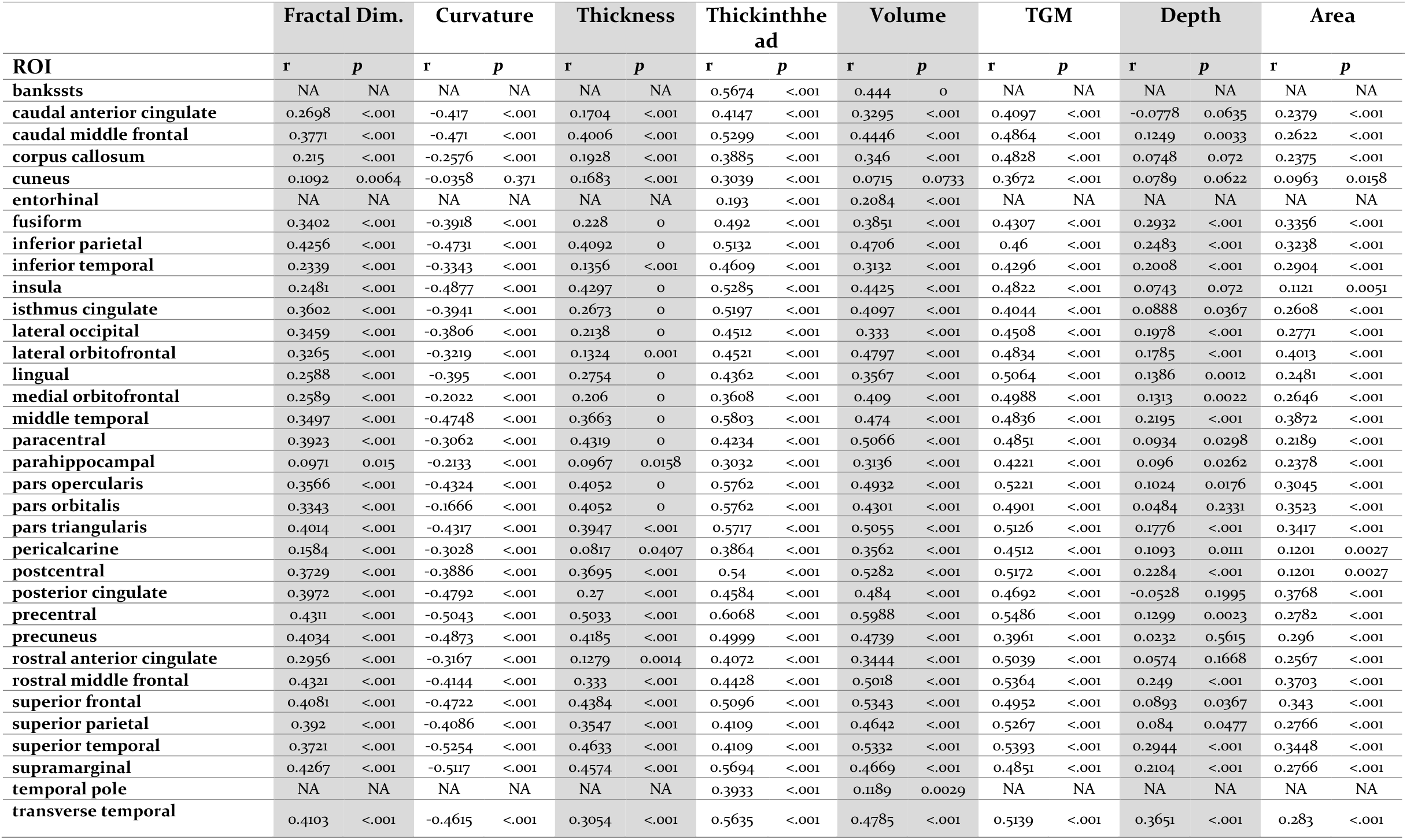
Regional fluid intelligence correlations in Cam-CAN. All p-values are FDR corrected at alpha = 0.05.

**Table 7:**
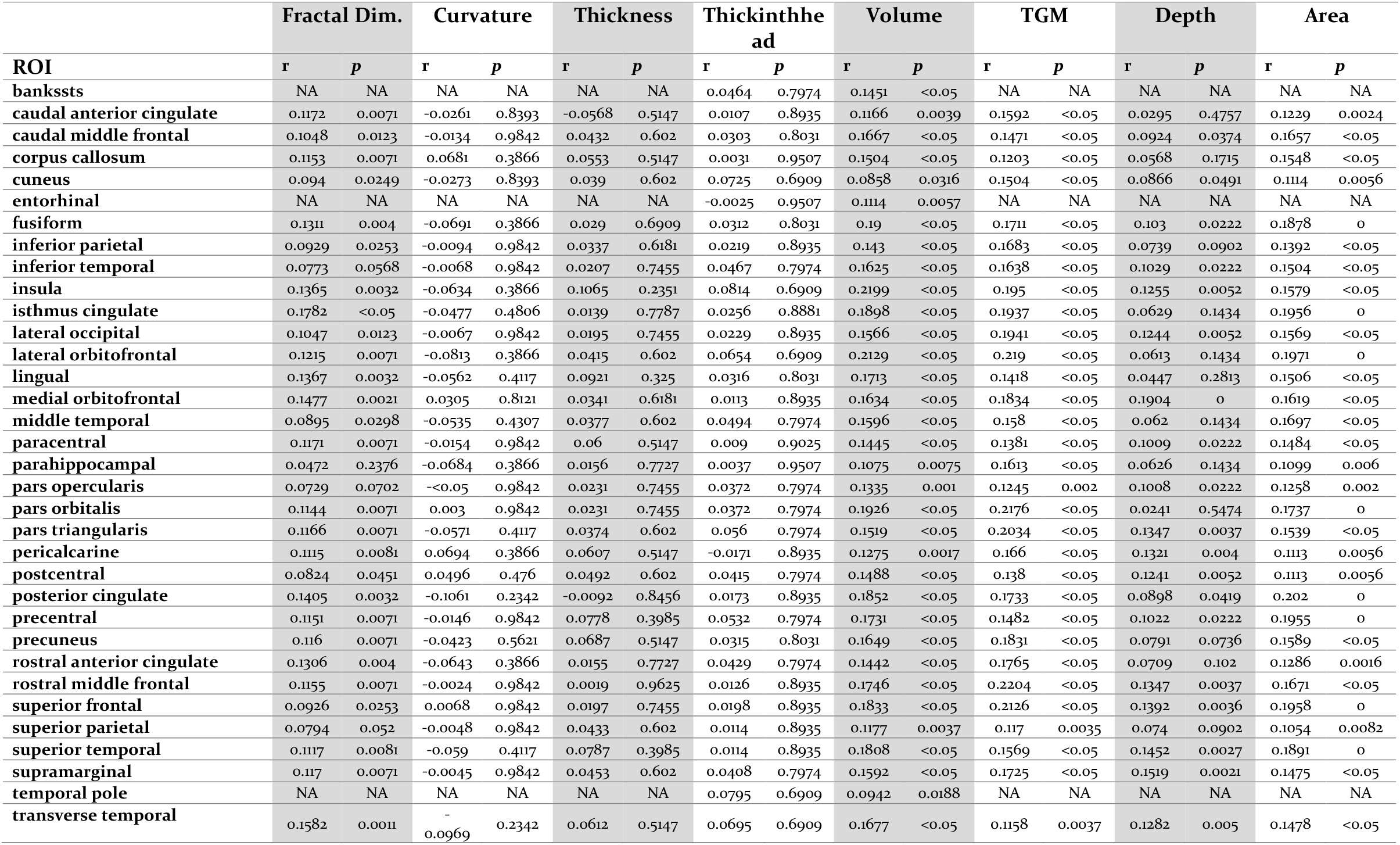
Regional age-residualized fluid intelligence correlations in Cam-CAN. All p-values are FDR corrected at alpha = 0.05.

**Figure 9:**
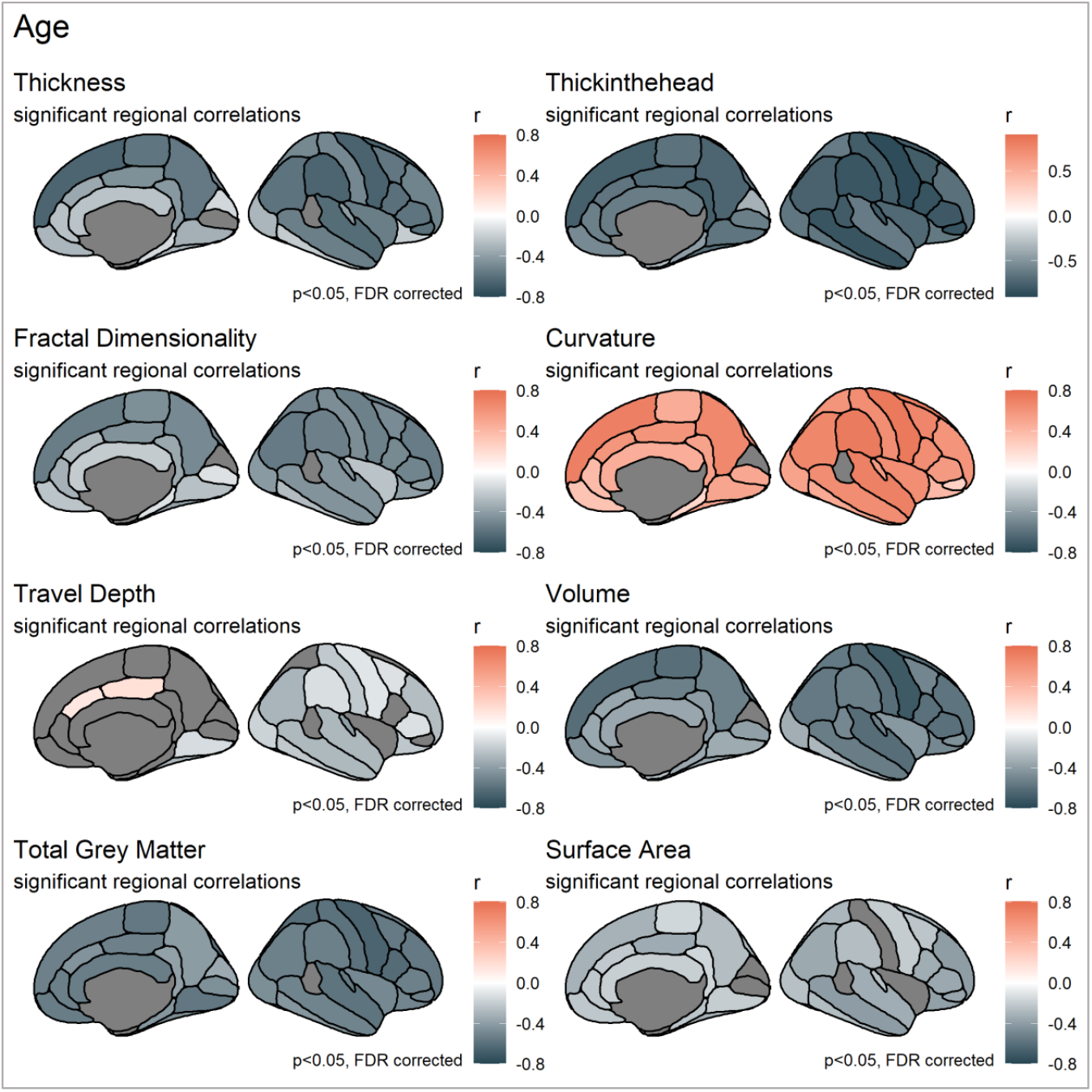
Significant regional age correlation for each metric. FDR corrected at alpha = 0.05.

**Figure 10:**
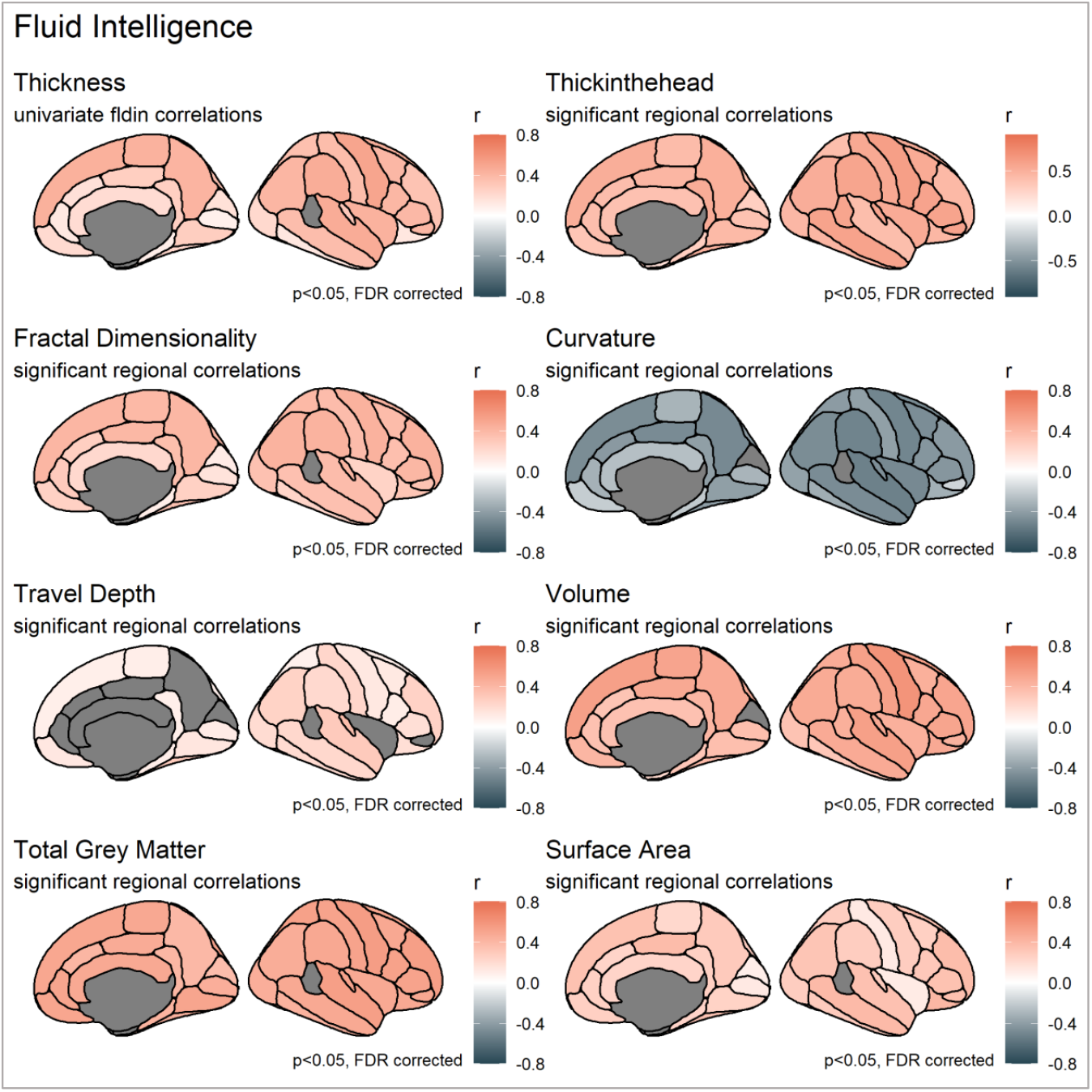
Significant regional fluid intelligence correlation for each metric. FDR corrected at alpha = 0.05.

**Figure 11:**
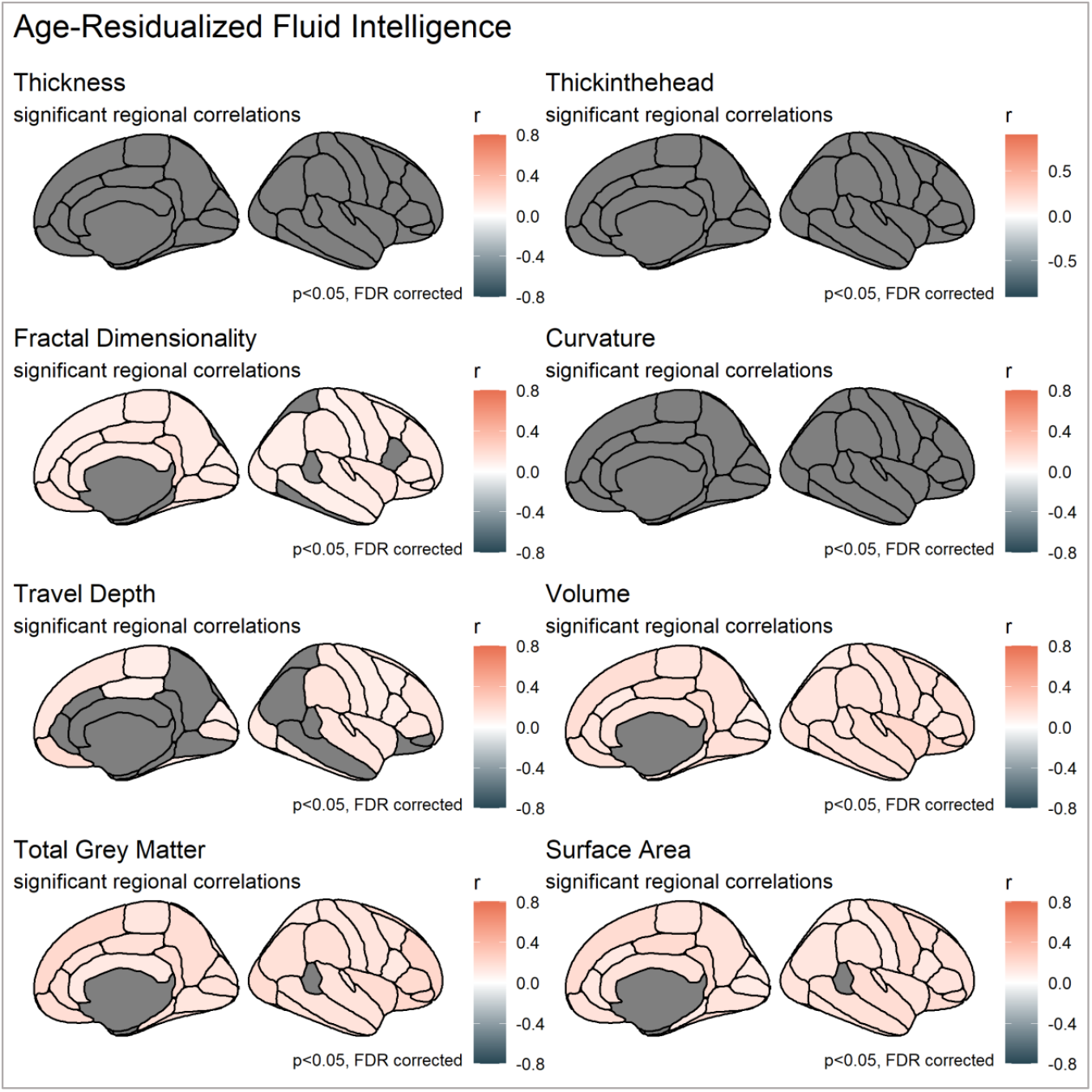
Significant regional age-residualized fluid intelligence correlation for each metric. FDR corrected at alpha = 0.05.

### 6 Longitudinal results

First, to assess whether Cattell test type (online versus pen/paper) made a difference, we tested for metric invariance and scalar invariance in the wave two cognitive data. This led to negligible drops in model fit (ΔCFI = 0.008 and 0.004 for metric and scalar invariance, respectively, Cheung & Rensvold, 2002), suggesting that assuming pencil and paper vs computer-based testing had equal measurement properties did not adversely affect the measurement of fluid intelligence. For all further analysis, this grouping factor was therefore ignored. Second,to ensure comparability of cognitive scores across Time 1 and Time 2, we tested for longitudinal measurement invariance (Widaman, Ferrer & Conger, 2010). We found that imposing invariance did not meaningfully decrease model fit (ΔCFI = 0.002; Cheung & Rensvold, 2002), suggesting longitudinal measurement invariance is tenable, and we were able to proceed to interpret change scores in the latent factor. Following the above inspections, we used Latent Change Score Models (LCSM) to examine morphometric and cognitive change over time.

**Table 8:** Cam-CAN raw scores and descriptive statistics for age, Cattell and longitudinal brain structure metrics

|  | Time | N | Mean | Minimum | Maximum | SD | Skewness | Excess kurtosis |
| --- | --- | --- | --- | --- | --- | --- | --- | --- |
| **Age** | T1 | 261 | 54.97 | 19.25 | 89 | 18.17 | -0.02 | -1.16 |
|  | T2 | 261 | 56.32 | 21.25 | 91.58 | 18.2 | -0.03 | -1.18 |
| **Cattell (sum score)** | T1 | 215 | 32.50 | 12 | 44 | 6.06 | -0.39 | -0.10 |
|  | T2 | 215 | 30.42 | 10 | 44 | 6.65 | -0.76 | 0.80 |
| **Surface Area** | T1 | 261 | 2527.43 | 1896.25 | 3299.01 | 256.81 | 0.22 | -0.22 |
|  | T2 | 261 | 2521.75 | 1898.46 | 3297.51 | 255.73 | 0.23 | -0.21 |
| **Cortical Thickness** | T1 | 261 | 2.61 | 2.28 | 2.89 | 0.1 | -0.19 | 0.45 |
|  | T2 | 261 | 2.6 | 2.29 | 2.91 | 0.1 | -0.19 | 0.3 |
| **Volume** | T1 | 261 | 7175.41 | 5417.15 | 9412.12 | 822.25 | 0.44 | -0.05 |
|  | T2 | 261 | 7124.88 | 5342.85 | 9311.37 | 824.73 | 0.42 | -0.05 |

**Table 9:** Second order latent change score model fit indices Cam-CAN.

| Cam-CAN |  | Model Fit Indices |  |  |  |  |  |
| --- | --- | --- | --- | --- | --- | --- | --- |
| Metric | Model | $\chi^2$ | $p$ | RMSEA [90 % CI] | CFI | SRMR | Yuan-Bentler scaling factor |
| **Thickness** | **FIML** | 5.275 | 0.072 | 0.039 [0.000, 0.072] | 0.992 | 0.026 | 0.763 |
| **Surface Area** | **FIML** | 4.228 | 0.121 | 0.033, [0.000, 0.079] | 0.997 | 0.015 | 0.721 |
| **Volume** | **FIML** | 3.655 | 0.161 | 0.028 [0.000, 0.065] | 0.995 | 0.014 | 1.468 |

**Table 10:** Second order latent change score model fit indices LCBC.

| Model Fit Indices |  |  |  |  |  |  |
| --- | --- | --- | --- | --- | --- | --- |
| Metric | $\chi^2$ | *p* | RMSEA [90 % CI] | CFI | SRMR | Yuan-Bentler scaling factor |
| **Thickness** | 13.605 | 0.001 | 0.090 [0.050, 0.135] | 0.993 | 0.038 | 1.070 |
| **Surface Area** | 2.418 | 0.298 | 0.033, [0.000, 0.079] | 0.999 | 0.007 | 1.091 |
| **Volume** | 47.648 | 0.000 | 0.178 [0.133, 0.227] | 0.975 | 0.034 | 0.845 |

**Figure 12:**
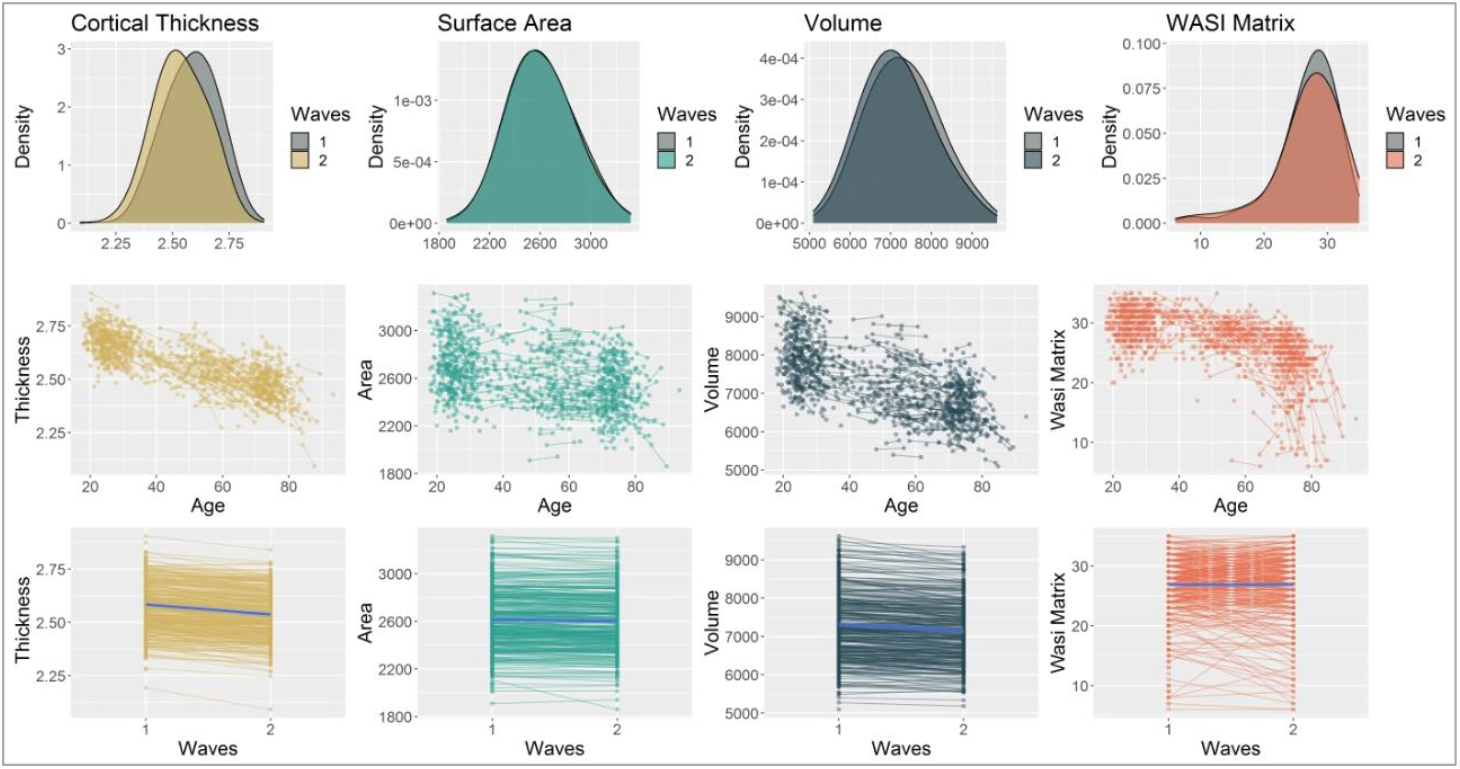
changes in volume, cortical thickness, surface area and fluid intelligence between time point 1 and time point 2 in LCBC sample.

**Figure 13:**
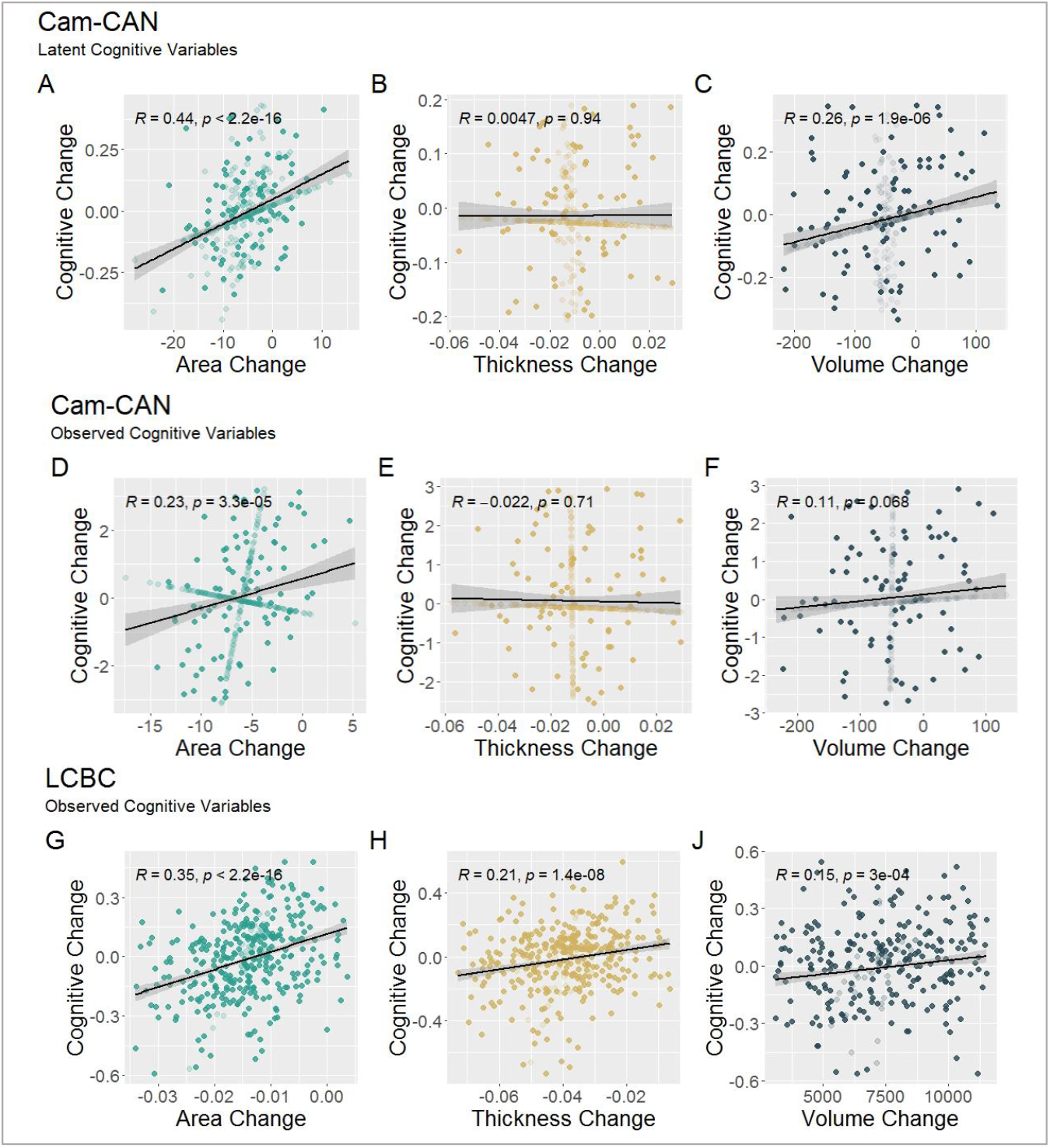
correlations of cognitive change and neural change in Cam-CAN (A-F) and LCBC (G-J). Shows that change in surface area is most strongly associated with cognitive change. Models A-C include latent cognitive variables, which were not possible to derive from the LCBC data, where we used observed cognitive scores instead. To compare like-for-like models, we include Cam-CAN observed variable models here, too (D-F). Note that the shaded dots are the models’ missingness estimates.

## Power Analyses, Morphometric Double Dissociation

*Sophia Borgeest*

*23/09/2021*

### Intro

Here we use R’s pwr package to run power analyses on the brain-age and brain-cognition relationship for volume, thickness and surface area. These include estimated correlation coeffcients, based on well-powered findings in the literature.

### Age

First, let’s run power analyses based on whole brain-age effect sizes (correlation coeffcients) found in the literature. We use this well-powered study as a source of reference (see Table 1 for whole brain - age correlation coeffcients):

https://www.sciencedirect.com/science/article/pii/S0197458010003210?casa_token=lUY7YAgJKZsAAAAA:FCrWz1X7EWi5lKjsFmzGBYMzKnVknQ8_X2iBUn3xqqdd-R3wU1pPnHEOasgn0XUZ175R4JtpXdvV

As a reminder, CamCAN has a sample size of N = 647, LCBC has N = 1345.

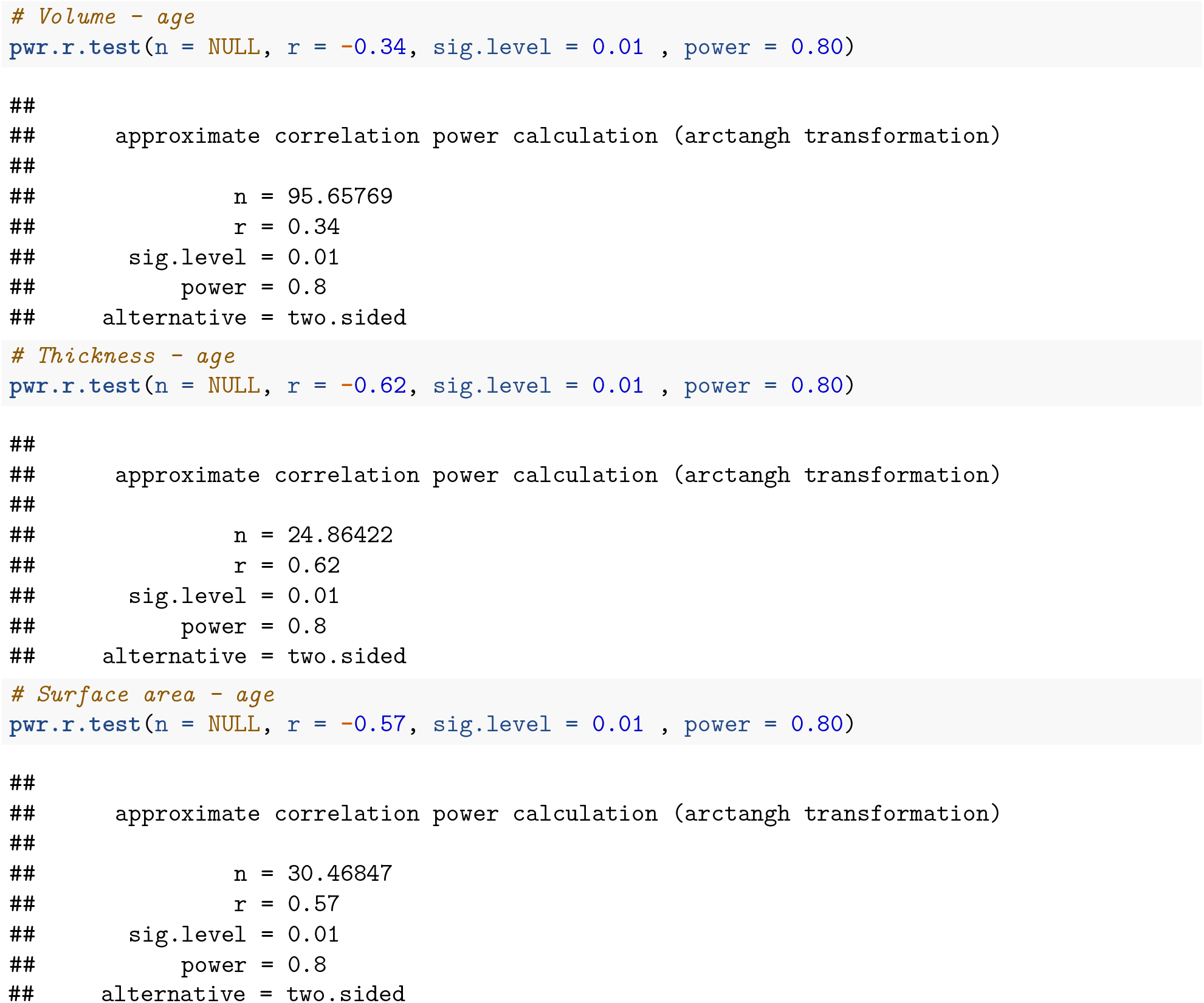

### Fluid intelligence

For volume and thickness, we use correlation coeffcients from this study (see Figure 3): https://www.sciencedirect.com/science/article/pii/S105381192030063X

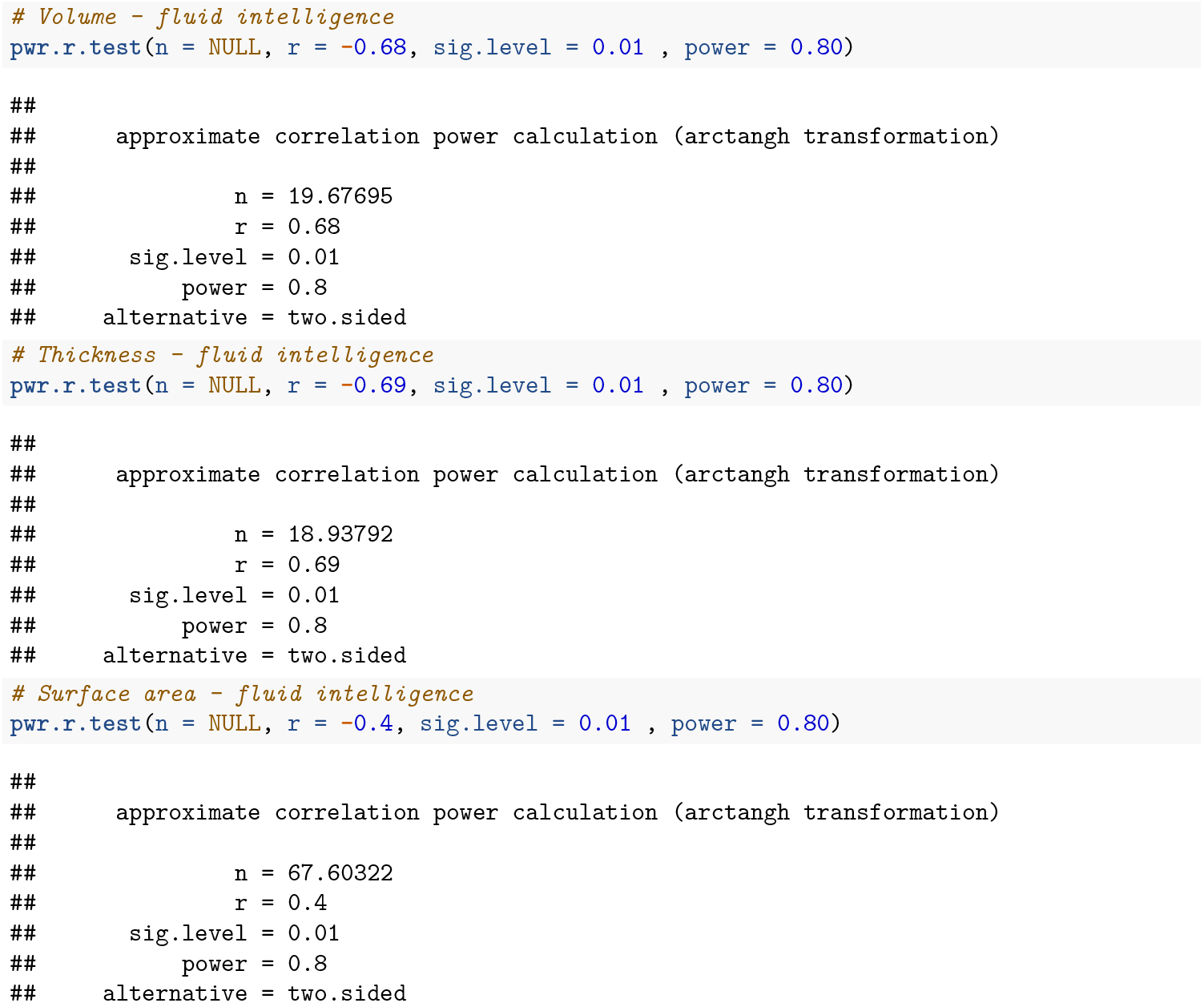

### Age-residualized fluid intelligence

Because very few studies have age-residualized cognitive abilities, no reliable, well-powered correlation coeffcients were availble in the literature. We therefore did not run a priori power analyses for these correlations.

